# Hybrid immunity to SARS-CoV-2 arises from serological recall of IgG antibodies distinctly imprinted by infection or vaccination

**DOI:** 10.1101/2024.01.22.576742

**Authors:** William N. Voss, Michael A. Mallory, Patrick O. Byrne, Jeffrey M. Marchioni, Sean A. Knudson, John M. Powers, Sarah R. Leist, Bernadeta Dadonaite, Douglas R. Townsend, Jessica Kain, Yimin Huang, Ed Satterwhite, Izabella N. Castillo, Melissa Mattocks, Chelsea Paresi, Jennifer E. Munt, Trevor Scobey, Allison Seeger, Lakshmanane Premkumar, Jesse D. Bloom, George Georgiou, Jason S. McLellan, Ralph S. Baric, Jason J. Lavinder, Gregory C. Ippolito

**Affiliations:** Department of Molecular Biosciences, The University of Texas at Austin, Austin, TX, USA; Department of Epidemiology, University of North Carolina at Chapel Hill, Chapel Hill, NC, USA; GSK Vaccines Institute for Global Health, 53100 Siena, Tuscany, Italy; Department of Chemical Engineering, The University of Texas at Austin, Austin, TX, USA; Basic Sciences Division and Computational Biology Program, Fred Hutchinson Cancer Center, Seattle, WA, USA; Department of Microbiology and Immunology, University of North Carolina at Chapel Hill, Chapel Hill, NC, USA; Department of Chemistry, The University of Texas at Austin, Austin, TX, USA; Howard Hughes Medical Institute, Seattle, WA USA

**Author notes:** These authors contributed equally (William N. Voss, Michael A. Mallory). Correspondence (J.J.L.), (G.C.I.).

**Keywords:** COVID-19, SARS-CoV-2, plasma, broadly neutralizing monoclonal antibody (bNAb), Ig-Seq, Cryo-EM, hybrid immunity, antibody feedback, immunological imprinting

## Abstract

We used plasma IgG proteomics to study the molecular composition and temporal durability of polyclonal IgG antibodies triggered by ancestral SARS-CoV-2 infection, vaccination, or their combination ("hybrid immunity"). Infection, whether primary or post-vaccination, mainly triggered an anti-spike antibody response to the S2 domain, while vaccination predominantly induced anti-RBD antibodies. Immunological imprinting persisted after a secondary (hybrid) exposure, with >60% of the ensuing serological response originating from the initial antibodies generated during the first exposure. We highlight one instance where hybrid immunity arising from breakthrough infection resulted in a marked increase in the breadth and affinity of a highly abundant vaccination-elicited plasma IgG antibody, SC27. With an intrinsic binding affinity surpassing a theoretical maximum (K_D_ < 5 pM), SC27 demonstrated potent neutralization of various SARS-CoV-2 variants and SARS-like zoonotic viruses (IC_50_ ∼0.1–1.75 nM) and provided robust protection *in vivo*. Cryo-EM structural analysis unveiled that SC27 binds to the RBD class 1/4 epitope, with both VH and VL significantly contributing to the binding interface. These findings suggest that exceptionally broad and potent antibodies can be prevalent in plasma and can largely dictate the nature of serological neutralization.

**HIGHLIGHTS:**

▪ Infection and vaccination elicit unique IgG antibody profiles at the molecular level
▪ Immunological imprinting varies between infection (S2/NTD) and vaccination (RBD)
▪ Hybrid immunity maintains the imprint of first infection or first vaccination
▪ Hybrid immune IgG plasma mAbs have superior neutralization potency and breadth

## INTRODUCTION

The COVID-19 pandemic is defined by the abrupt emergence, periodic cycles of rapid evolution, and subsequent extinction of ancestral SARS-CoV-2 variants of concern (VOC), primarily in response to the ongoing development of expanding immunity.^1^ The immunoglobulin G (IgG) antibody response against the SARS-CoV-2 spike (S) attachment glycoprotein (IgG anti-S) is an essential component of humoral immunity and its quantity (or titer) in plasma has been correlated with protection from COVID-19 disease.^2^ Primary humoral immunity elicited by infection or vaccination results in diversified antibody repertoires capable of recognizing multiple S epitopes^3–6^ which mature over time and potentiate high-affinity and cross-neutralizing responses to SARS-CoV-2 VOCs^7–9^. Subsequently, memory B cells (MBCs) and their antibody-secreting progeny, plasmablasts (PBs), also contribute to humoral immunity through the rapid production of IgG antibody during recall responses to a secondary challenge with the same or related virus. Furthermore, immune recall improves the bulk quality of plasma IgG breadth, potency, and durability to SARS-CoV-2 variants.^9,10^ Despite these significant observations, our comprehension of SARS-CoV-2 humoral immunity remains fragmentary at a molecular level, particularly the IgG proteome and its constituents which comprise anti-S plasma repertoires.

The quality (breadth, potency, and durability) of protection conferred by humoral immunological memory is intimately linked to the clonal diversity (relative abundance and polarization of the constituent lineages) and spectrum of epitopes targeted by a polyclonal IgG plasma repertoire. Obstacles hindering the study of polyclonal IgG repertoires may include overwhelming diversity, the temporal waning of antigen-specific titers, and *immunological imprinting*^11,12^—first described in 1956 as *persistent antibody orientation*^13^— which refers generally to the fixating effects that original exposures have on subsequent antibody responses to antigenically drifted but related viruses. In turn, future waves of B cell activation may be subject to regulatory control mechanisms like IgG antibody feedback^14–16^ wherein pre-existing antibodies inhibit B cell (re)activation and new antibody formation. Furthermore, an increasingly significant portion of the human population has acquired *hybrid immunity* to SARS-CoV-2, stemming from the combined protection of natural immunity following repeated cycles of infection and vaccination. While the profound impacts of immunological imprinting resulting from prior exposure to SARS-CoV-2 and its various effects on hybrid immune enhancement and immune recall have been documented^8,17–20^, the phenomenon remains unresolved at the molecular level of the IgG anti-S plasma proteome and its individual components. While it is widely acknowledged that hybrid priming through infection and vaccination generally enhances immunity^19,21–23^, there is a need for additional clarity regarding the imprint left by the ancestral virus infection or vaccination at the pandemic’s onset. This includes understanding the impact of the chronological sequence of infection preceding vaccination on the serological IgG repertoire, or vice versa, and the subsequent patterns established by early VOCs.

Here we present a comparative molecular-level investigation of the polyclonal plasma IgG response to the SARS-CoV-2 S glycoprotein, whether elicited by natural infection (primary ancestral SARS-CoV-2 virus or pre-Omicron VOC breakthrough infection), vaccination (stabilized ancestral Wuhan-Hu-1 spike), or their combination. Relative quantitation demonstrates that the primary IgG response is imprinted predominantly against epitopes residing outside the receptor-binding domain (RBD) upon infection, but RBD epitopes upon vaccination. Moreover, secondary plasma IgG recall responses are heavily predetermined by this initial imprint and derive from the original primary lineages. The combined impact of infection and vaccination results in hybrid immunity that can exploit a broad and potent neutralization epitope conserved in SARS-related coronaviruses^24,25^ (“class 1/4”^26,27^), exemplified in our study by the RBD-targeting monoclonal antibody (mAb) SC27. This mAb displayed an intrinsic binding affinity of its monovalent antigen-binding fragment (F_ab_) surpassing the reported K_D_ of all other known human antibodies in published research, rivaling the breadth and potency of erstwhile FDA-approved mAbs. SC27 potently neutralizes ancestral and contemporary SARS-CoV-2 VOC and many antigenically distinct zoonotic sarbecoviruses that are poised for human emergence.

## RESULTS

### IgG anti-S binding profiles differ at the bulk level between infection and vaccination and imprint upon distinct S domains

For ancestral SARS-CoV-2 primary infection, we established previously that only a minor fraction (<25%) of the circulating S-binding IgG antibody repertoire targets the receptor-binding domain (RBD), whereas the majority of IgG lineages (>75%) are directed against the N-terminal domain (NTD) or the conserved S2 subunit.^6^ Motivated by earlier findings based upon bulk serological responses,^28,29^ we posited that this distinctive prevalence of non-RBD targeting in primary infection might carry over to subsequent vaccinations, as opposed to those vaccinated without prior infection (naïve vaccinees). To examine this hypothesis, we assessed ratiometric S glycoprotein domain specificity using an RBD competition ELISA in an early pandemic cohort (n=13) and observed that immunological imprinting through ancestral S exposure differs between infection (with augmentation towards S2 and NTD) and vaccination (with augmentation towards RBD), and this differential antibody orientation persists in hybrid immune individuals (**Figure 1A**).

**Figure 1.**
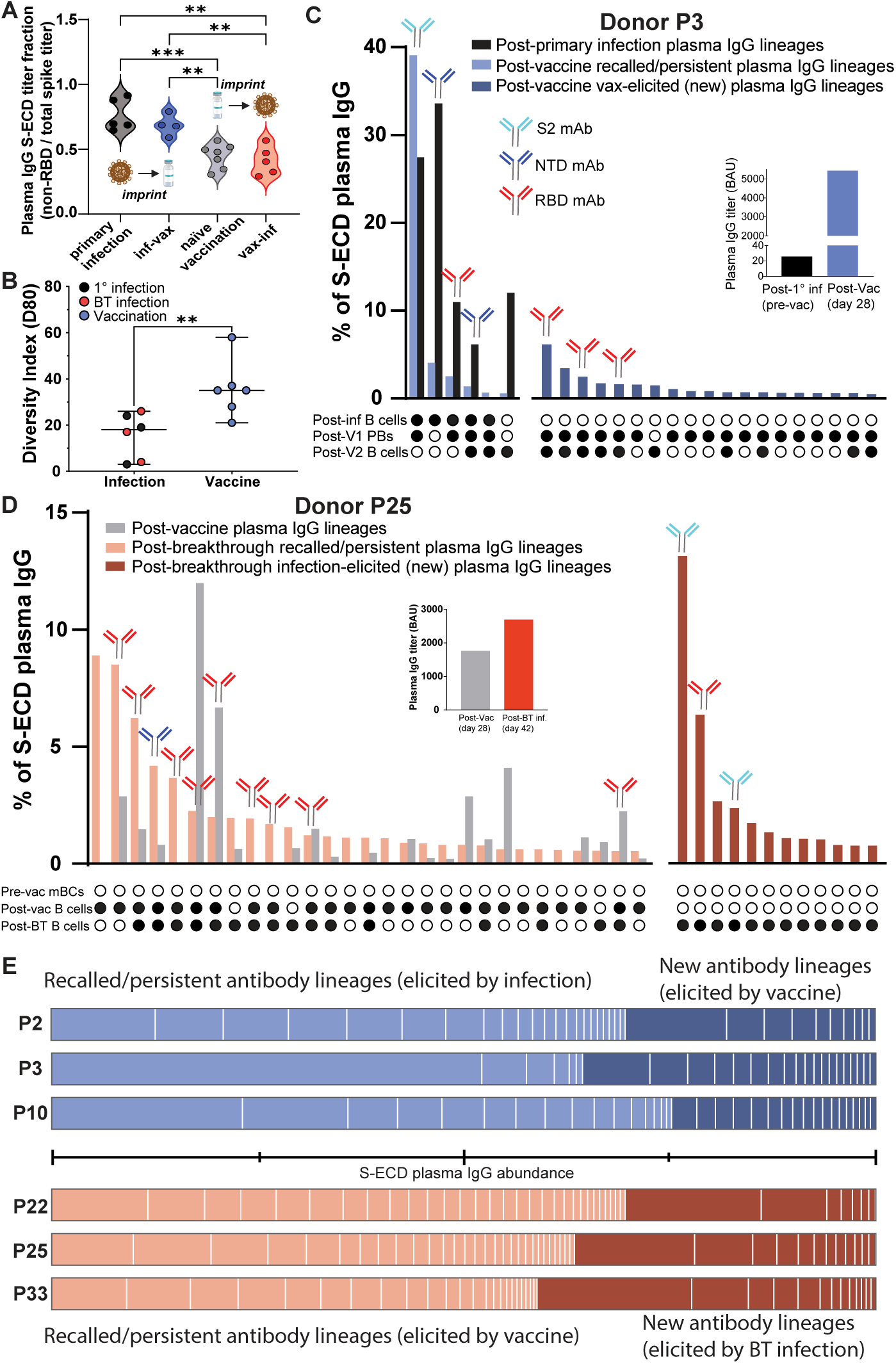
IgG serological recall and hybrid immunity are predetermined by the initial immunological imprint set by infection or vaccination. (A) Plasma RBD competition ELISA reveals that mode and order of exposure to SARS-CoV-2 spike results in differential persistent antibody orientation, with infection and vaccination imprinting non-RBD and RBD epitopes, respectively. (B) Vaccination-induced plasma IgG repertoires are significantly more diverse than those induced by infection. (C) Donor P3 plasma IgG repertoire elicited by primary infection (light blue bars), recalled by subsequent vaccination (black bars), and newly elicited by subsequent vaccination (dark blue bars). Each bar represents an individual plasma IgG lineage. Antibody symbols above bars indicate spike domain specificity of recombinantly cloned mAbs representative of each lineage. In both (C) and (D), the inserts show anti-spike plasma binding titers at each time point, and the upset plots below the repertoire bar plots indicate whether the plasma lineage was detected (filled circle) in total B cells, sorted MBCs, and sorted PBs. “Post-V1” = following the first vaccine dose, and “Post-V2” = following the second vaccine dose. (D) Donor P25 plasma IgG repertoire elicited by naïve vaccination (pink bars), recalled by subsequent infection (grey bars), and newly elicited by subsequent vaccination (red bars). Insert shows anti-spike plasma binding titers at each time point. (E) Hybrid immunity: across the cohort, most plasma IgG lineages following secondary exposure are recalled from lineages originally imprinted by the initial exposure. The horizontal line between the plots denotes each quartile of the plasma IgG repertoire by relative abundance. Significant differences calculated using the Mann-Whitney *U* test linked by horizontal lines are indicated by asterisks: **p < 0.01, ***p < 0.001.

### IgG anti-S neutralization profiles differ at the bulk level between infection and vaccination but are equilibrated by hybrid immunity

These observations raised the question whether bulk-level changes in antibody binding titer and differential S domain specificity may impact neutralization profiles and the functional composition of IgG anti-S proteomes. Within our early pandemic cohort, six individuals for whom we obtained longitudinal blood draws across multiple spike exposures **(Figure S1A)** were examined for a comprehensive analysis of the combination of SARS-CoV-2 infection, vaccination, or breakthrough (BT) infection. Three (P2, P3, P10) had a primary infection with ancestral virus during the first wave of COVID-19 in the U.S.A. in March or July 2020 and were vaccinated 9-13 months later (“infection–vaccination” convalescent group), whereas the three other donors (P22, P25, P33) were first immunized during the inaugural release of mRNA vaccines in January and February of 2021 and subsequently experienced a pre-Omicron BT infection in the U.S.A. 5-8 months later between July–October 2021 during the Delta VOC surge (“vaccination–infection” naïve group) **(Table S1)**.

First, we characterized bulk serology in the two groups by performing indirect IgG ELISA against recombinant S ectodomain (S-ECD) proteins (stabilized ancestral Wuhan-1 S-6P [HexaPro^30^] and Omicron BA.1) followed by live virus neutralization assays (ancestral WA1/2020 and Omicron BA.1) using plasma from each donor and time point (**Figure S1B**). For the infection–vaccination convalescent group, vaccination elicited a robust increase in binding and neutralization titer (NT_50_; reciprocal plasma dilution resulting in 50% neutralization titer) against ancestral S or virus, respectively, as expected for a recall response.^31,32^ For individuals in the vaccination–infection naïve group, vaccination elicited comparatively lower S-binding and virus neutralization titers, but subsequent BT infection by a pre-Omicron variant induced substantial increases in neutralization titers that were comparable to the convalescent group’s post-vaccination levels (**Figure S1B**). For both groups, the anti-S recall response boosted titers against not only the ancestral virus but also the prospective Omicron BA.1 VOC which had not yet emerged. In summary, and in agreement with previous reports^9,18,21,22,33^, our findings in a select cohort corroborated a significant enhancement in plasma IgG binding titer, breadth, and neutralization potency among individuals with hybrid immunity.

### IgG anti-S binding profiles and antibody diversity differ at the molecular level between infection and vaccination

Quantitative assessment of antibody levels using analytical methods like ELISA lacks the ability to distinguish abundances at the lineage (family) or clonal (individual) level. Thus, we sought to establish at the molecular level whether the bulk increases in binding and neutralization titers observed in the hybrid immune individuals stem from an augmentation in pre-existing plasma antibody lineages, or if newly elicited antibody lineages with novel specificities are the determining factor. The lineage composition and relative abundance of IgG antibodies comprising the polyclonal plasma response to stabilized HexaPro S-ECD were determined using the Ig-Seq pipeline^34–37^ that integrates LC–MS/MS proteomics of chromatographically enriched antigen-reactive polyclonal IgG with high-throughput sequencing of B-cell heavy-chain (VH), light-chain (VL), and single B-cell VH:VL variable region repertoires (BCR-Seq).

Overall, the repertoire diversity index (D80, **Figure 1B**) of anti-S-ECD plasma IgG lineages varied significantly between post-infection and post-vaccination (*p* < 0.01). Infection, whether primary or breakthrough, resulted in more polarized (i.e., top-heavy in terms of relative abundance of IgG lineages) plasma IgG repertoires with an average D80 of 17 plasma IgG lineages, whereas vaccination resulted in >2-fold greater diversity (avg D80 = 36 plasma IgG lineages) against the viral spike protein.

Primary infection in one individual, donor P3, induced a strikingly restricted pauciclonal response comprising only six IgG anti-S lineages (D80 = 4) with the two most abundant ones accounting for >60% of the total (directed towards the NTD and S2 domains, respectively) (**Figure 1C**). All six IgG lineages in the post-infection plasma were later detected after mRNA vaccination, which significantly amplified these lineages, as evidenced by the >100-fold rise in plasma IgG anti-S titer determined by ELISA (**Figure 1C**, inset). This amplified response was underscored at the B-cell level by the facile detection of clonally related plasmablasts (PBs) at day 7 post-1st-dose vaccination for 4 of the 6 lineages (**Figure 1C**). Vaccination not only boosted these 6 pre-existing lineages but markedly diversified the IgG anti-S repertoire and elicited 19 new IgG lineages in convalescent donor P3 (D80 = 37). Of the top 5 vaccine-elicited lineages, three were produced as recombinant mAbs and, notably, all three were specific to the RBD (**Figure 1C**).

Polyclonal diversification of the IgG anti-S repertoire by mRNA vaccination was also observed in a naïve individual, donor P25 (D80 = 35; **Figure 1D**). We expressed and tested 12 of the vaccine-induced lineages as recombinant mAbs, which collectively comprised 48% of the IgG anti-S repertoire **(Figure S2)**. Like donor P3, the vaccine-induced lineages in this donor were prevailingly oriented towards the RBD, as nine of the twelve were RBD reactive (**Figure S2**). After BT infection, we detected the emergence of 12 new lineages (**Figure 1D**). Two of these newly identified BT lineages, which represented a significant share of the total repertoire by rank and abundance (rank 1: 13.1%; rank 4: 2.4%), recognized the S2 subunit, reflecting the non-RBD S-domain bias we consistently observe in SARS-CoV-2 infections.

**Figure 2.**
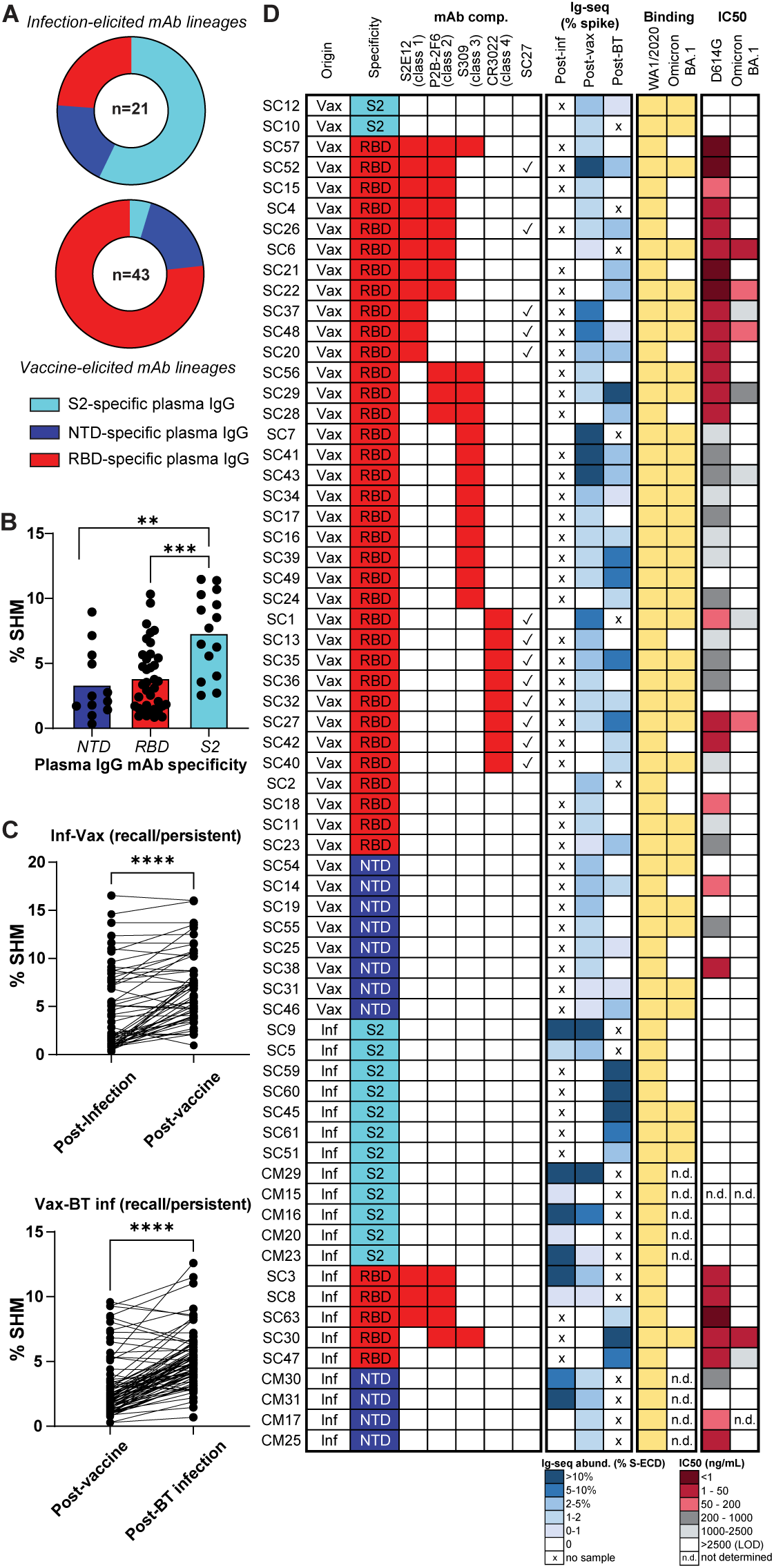
Molecular IgG responses: Differential immunological imprinting by infection (S2/NTD) and vaccination (RBD). (A) Representative mAbs from infection-elicited lineages mostly bind non-RBD epitopes (top) whereas those from vaccine-elicited lineages tend to bind RBD epitopes (bottom). (B) Mean VH somatic hypermutation rates are highest in S2-directed lineages, followed by RBD- and then NTD-directed lineages. (C) VH sequences of recalled plasma IgG lineages are significantly more mutated after the second spike exposure. (D) Heatmap displaying the origin, spike domain specificity, RBD class (when applicable), relative abundance after infection and/or vaccination, binding to Wuhan-Hu-1 and Omicron BA.1, and *in vitro* neutralization capacity of the representative mAbs cloned and characterized from the donors in this study. “Origin” column indicates the time point at which the lineage was first detected (BCR-Seq or Ig-Seq). Significant differences calculated using the Mann-Whitney *U* test linked by horizontal lines are indicated by asterisks: **p < 0.01, ***p < 0.001, ****p < 0.0001.

In contrast to the observed pattern in individuals who experienced prior infection followed by vaccine-induced antibody recall (P2, P3, P10), individuals who were naïve vaccinees (P22, P25, P33) did not boost the majority of the highly prevalent pre-existing anti-S plasma IgG lineages upon recall by a subsequent challenge (i.e., BT infection) (**Figure S2**). Instead, BT infection selectively reactivated only a minor fraction of the vaccine-elicited plasma IgG repertoire, constituting <20% of lineages, in the case of the three naïve vaccinees.

### Hybrid immunity is predetermined by recall of the initial immunological imprint set by infection or vaccination

Across all six donors, most plasma IgG lineages detected after a subsequent challenge are those originally imprinted by the initial exposure (**Figure 1E**). Within the infection–vaccination group, a significant proportion of the post-vaccine plasma IgG abundance, ranging from 64% to 75%, is imprinted by the initial primary infection. Similarly, within the vaccination–infection group, a comparable range of 59% to 70% of the post-BT infection plasma IgG abundance is imprinted by the vaccine. Together, these molecular-level data illustrate how the polyclonal IgG serological recall of durable antibody lineages is dictated by the initial immunological imprint established by infection or vaccination with ancestral spike.

### Immunological imprinting by infection (S2/NTD) versus vaccination (RBD) resolved at monoclonal resolution

A total of 66 plasma IgG mAbs identified across the six donors (and circulating at >0.5% relative abundance) were expressed recombinantly to determine their S-domain specificity and functionality. Among the 21 plasma IgG lineages induced by infection (primary or BT), most of their corresponding mAbs (16 out of 21) bound to regions outside of the RBD (**Figure 2A**), in agreement with our bulk serological findings (**Figure 1A**), indicating a predominantly non-RBD response to infection. In contrast, plasma IgG lineages induced by vaccination (n=43) preferentially targeted the RBD (33 out of 43, **Figure 2A**), consistent with bulk serology data.^28^

Among the infection-elicited plasma IgG mAbs that bound specifically to the S2 domain, 7 of 12 mapped to highly abundant lineages (>10% by relative abundance) (**Figure S2**, **Figure 1C**, **Figure 1D**). These anti-S2 mAbs had significantly higher somatic hypermutation (SHM) levels compared to those specific for the RBD or NTD (**Figure 2B**; *p* = 0.0004, *p* = 0.0014), suggestive of cross-reactive B-cell memory responses directed towards conserved epitopes shared between a previously encountered seasonal human coronavirus (HCoV) and SARS-CoV-2. All antibody lineages recalled by subsequent exposure to S antigen or BT virus, irrespective of domain specificity, were significantly more mutated than their progenitors (**Figure 2C**; *p* < 0.0001).

To ascertain functionality, we examined the plasma IgG mAbs by indirect ELISA and live virus neutralization assays against the ancestral (WA1/2020 D614G) and Omicron (BA.1) spike protein and viruses (**Figure 2D**). None of the S2-directed mAbs were capable of neutralization. Out of the 12 NTD-directed mAbs, however, six were able to neutralize the D614G founder virus but lost neutralization against Omicron. As previously demonstrated by our laboratory and other research teams^6,38,39^, the NTD of the ancestral Wuhan-Hu-1 S protein contains a neutralization “supersite” epitope, which was lost in the Omicron VOC. In contrast to S2-directed and NTD-directed mAbs, a greater proportion of the RBD-directed mAbs were neutralizing, with 37/40 demonstrating neutralization half-maximal inhibitory concentration (IC_50_) <2500 ng/mL against ancestral D614G virus.

To determine if the epitope classes of the RBD-directed mAbs identified in this study were correlative with loss of neutralization against Omicron, we examined all 40 mAbs by competitive ELISA against control class 1-4 neutralizing RBD mAbs (S2E12,^40^ P2B-2F6,^41^ S309,^42^ and CR3022^43^) (**Figure 2D**). Based on structural studies^26,27^ and deep mutational scanning (DMS) approaches defining escape pathways^44^, RBD-targeting mAbs have been categorized into several classes (class 1 to 4), according to contact residues and binding to the up- or down-conformation of the RBD, as well as within or outside the receptor-binding motif (RBM). Thirty-five of the 40 mAbs binned with at least one of the control antibodies examined, with the largest bin (10/35) represented by combined class 1/2 competitive mAbs, along with the control mAbs S2E12 (class 1) and P2B-2F6 (class 2). Twenty of the 35 mAbs were competitive with a single control (n=3 class 1, n=9 class 3, and n=8 class 4). The ability to cross-neutralize Omicron BA.1 was not restricted to any class 1-4 epitope bin. Interestingly, although the class 4 “cryptic epitope”^45^ is distal to the ACE2 binding site and is exposed only when the RBD is in the “up” position,^26^ two of the eight class 4 RBD mAbs were cross-neutralizing against Omicron (SC1 and SC27). Upon closer examination, we found that SC27 competed with all eleven class 1 and class 4 mAbs in our panel, indicating that SC27 at least partially sterically blocks class 1 antibodies as well.

### Hybrid immune IgG plasma mAbs have superior neutralization potency and breadth

Of the 33 plasma IgG RBD-directed mAbs derived from the vaccine–infection group (**Figure 2**), approximately half (n=18) were identified in the plasma of naïve vaccinees, and half (n=15) from BT infections. A comparison of the neutralization potency of these two groups of plasma IgG reveals the benefit of hybrid immunity in the blood plasma at a monoclonal resolution. With one exception, all the BT infection plasma mAbs have potent (<1 µg/mL) IC_50_ neutralization against the ancestral D614G virus, and the majority of these are ultra-potent (<50 ng/mL IC_50_) (**Figure 3A**). Further, when examining the ability of these plasma mAbs to cross-neutralize, 7 of 15 mAbs derived after BT infection show detectable (<2500 ng/mL LOD) neutralization of Omicron BA.1 (**Figure 3A**) whereas only 1 of 18 from the naïve vaccinees can neutralize this VOC. We conclude that BT infection-elicited RBD-directed antibodies are significantly enriched to target neutralizing spike epitopes, and often cross-react with Omicron VOCs, when compared to those generated by vaccination alone (**Figure 3A**; *p* < 0.01).

**Figure 3.**
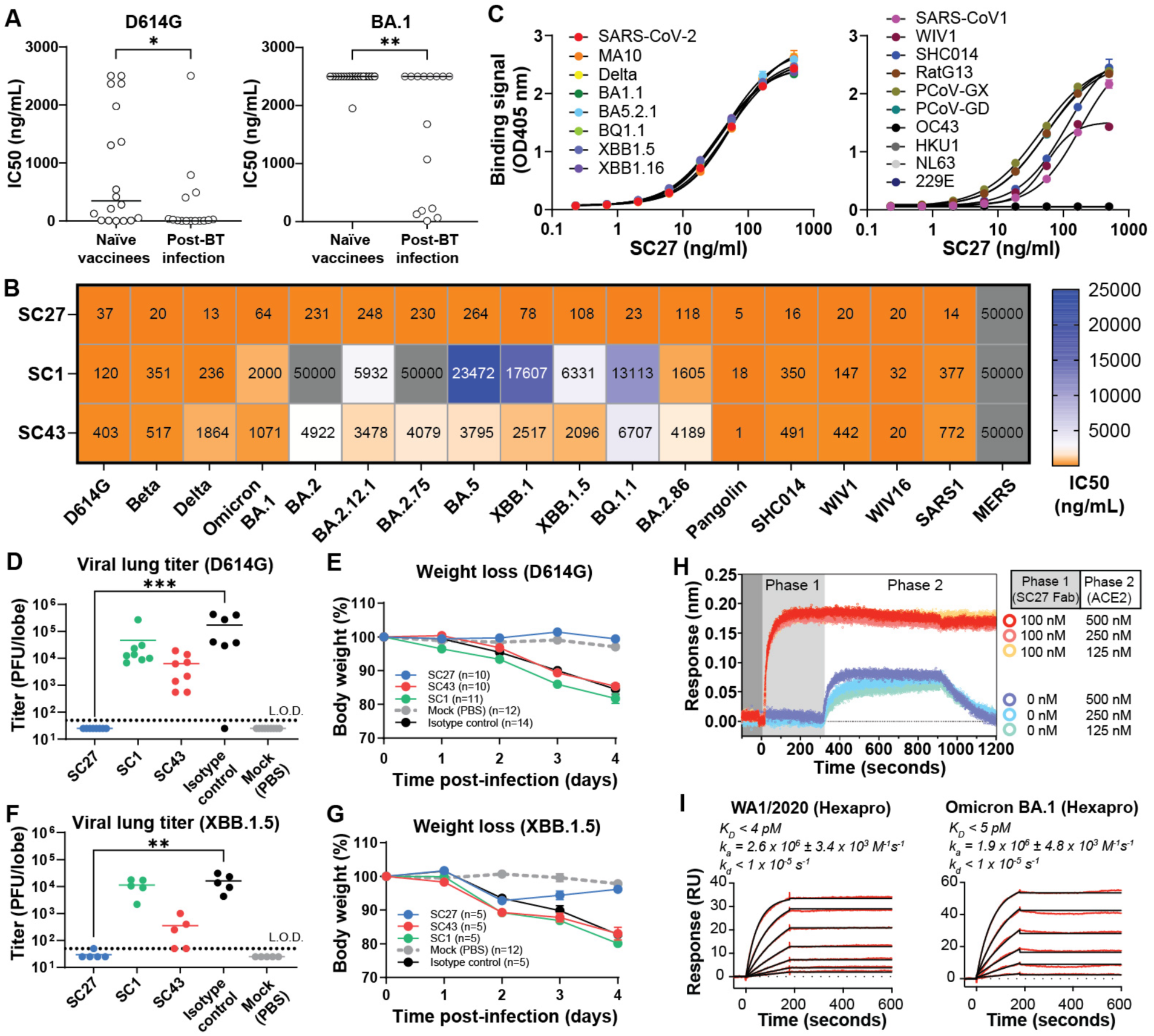
Identification of the broad, potent, and protective plasma mAb SC27. (A) Compared with naïve vaccination, BT infection elicits a larger fraction of RBD-directed antibodies endowed with greater neutralization potency (Wuhan D614G, left) and breadth (Omicron BA.1, right). (B) Live-virus neutralization assays using RBD-directed plasma mAbs screened against a wide panel of SARS-CoV-2 VOCs as well as zoonotic sarbecoviruses. (C) ELISA binding data to recombinant RBD proteins showing SC27 cross-reactivity across a broad panel of SARS-CoV-2 VOCs (left) and other sarbecoviruses (right). (D to G) *In vivo* prophylactic protection of 12-month-old BALB/c mice against standard intranasal infection challenge dose (10^3^ PFU) of mouse-adapted (MA10) SARS-CoV-2 (D614G or XBB.1.5). Error bars represent SEM about the mean. (H) Biolayer interferometry sensorgram demonstrating complete inhibition of SARS-CoV-2 Wuhan-Hu-1 spike–ACE2 binding by mAb SC27. (I) SPR sensorgram of SC27 Fab binding to stabilized (HexaPro) WA1/2020 and Omicron BA.1 spike ECD proteins. Significant differences calculated using the Mann-Whitney *U* test linked by horizontal lines are indicated by asterisks: *p < 0.05, **p < 0.01, ***p < 0.001.

### Ultra-high affinity and broadly neutralizing plasma mAb SC27: vaccine elicited, affinity matured by breakthrough infection, and protective *in vivo*

Next, we analyzed the neutralization capacity of cross-reacting plasma mAbs SC1, SC27, and SC43 (which fall within the class 4 and class 3 RBD epitope bins) against a larger panel of β-CoVs including the most recent VOCs (**Figure 3B**). These three plasma mAbs stood out in our panel because of their demonstrated ability to potently cross-neutralize divergent and pre-emergent nLUC-encoding zoonotic SARS-like CoVs, such as bat (SHC014, WIV1, WIV16)^46^ and pangolin (PCoV) live viruses,^47^ as well as SARS-CoV 2003,^48^ suggesting that they recognize conserved pan-sarbecovirus neutralizing epitopes. Although the progenitors for all three antibodies were elicited by mRNA vaccination, we conjectured that broad cross-neutralization is acquired only steadily over time with clonal evolution. This conjecture is neatly illustrated by antibody SC27 whose vaccine-elicited progenitor neutralizes ancestral WA1/2020 D614G, but not Omicron BA.1, whereas its BT infection counterpart potently neutralizes both (**Figure S3**). The SC27 VH region is encoded by the IGHV2-26 gene segment, which is rarely utilized by human B cells to generate RBD-directed antibodies following a primary S exposure,^49^ but its frequency increases substantially upon a hybrid immunological event as evidenced in our Ig-Seq dataset and as supported by published sequences deposited to the CoV-AbDab antibody database.^50^

SC27 neutralized authentic virus across a broad spectrum of early-generation SARS-CoV-2 VOCs (Beta, Delta), all first-generation Omicron sublineages of 2022 (BA.1, BA.2, BA.2.12.1, BA.2.75, BA.5, XBB.1, BQ.1.1), and even the most recent of Omicron variants in 2023 (XBB.1.5 and BA.2.86), with IC_50_ values ranging from 13–264 ng/mL (**Figure 3B**). Compared with the spike of BA.1, BA.2.86 possesses 29 additional mutations, including 14 mutations in the RBD. Beyond SARS-CoV-2, SC27 exhibited remarkable ultra-potency and not only neutralized SARS-CoV 2003 at 14 ng/mL but also effectively targeted zoonotic pangolin and bat CoVs within the range of 5–20 ng/mL. In line with its ability to neutralize a wide range of viruses, SC27 also exhibited cross-reactivity in a multiplex binding assay with a diverse array of CoV RBDs; however, this cross-reactivity was confined to the sarbecovirus lineage of β-CoVs and did not extend to the embecovirus (OC43, HKU1, NL63, 229E) or merbecovirus (MERS) lineages (**Figure 3C** and data not shown). We additionally tested SC27 for *in vivo* functional protection against SARS-CoV-2 infection (D614G and XBB.1.5) in the MA10 mouse model.^51^ SC27 demonstrates complete protection against viremia in the lungs and against wasting out to four days post-infection (**Figures 3D-G**). Consistent with its robust *in vitro* neutralization and *in vivo* protection, SC27 binding clashes with ACE2 binding to the RBD and completely ablates the ACE2–spike (D614G) interaction when tested by competition binding biolayer interferometry (BLI) (**Figure 3H**). The intrinsic binding affinity of SC27 F_ab_ surpasses the apparent K_D_ of all other human SARS-CoV-2 mAbs thus far reported, including former FDA-approved therapeutic antibodies.^52^ Surface plasmon resonance (SPR) binding analysis (**Figure 3I**) confirmed that SC27 Fab recognizes both ancestral (Wuhan-Hu-1) and Omicron variant (BA.1) S proteins with an exquisite intrinsic binding affinity of <5 pM, which falls beyond the realm of theoretical maxima for antibody association and dissociation rates (<10^6^ M^-1^ sec^-1^ and 10^-5^ sec^-1^, respectively) thought to be achievable via B-cell clonal selection and affinity maturation during endogenous immune responses.^53,54^

### Structural basis for SC27-mediated neutralization

To aid our understanding of the ultra-potent and broad neutralizing capacity of SC27, its structure in complex with Omicron BA.1 was determined by cryogenic electron microscopy (cryo-EM) at a global resolution of 2.6 Å (**Table S3, Figure S4**). The region near the RBD exhibited structural heterogeneity. Spike trimers were observed with either two or three SC27 Fabs bound, including a dimer-of-spike-trimers interface mediated by the RBDs (**Figures S5 and S6**). Extensive processing in 2D and 3D, followed by local refinement, improved the Coulomb potential map quality near the RBD and NTD (3.1 Å local resolution), resolving SC27 bound to the “inner face” of the RBD in the “up” conformation (**Figure S4**). The structure revealed a unique binding interface along the highly conserved class 4 cryptic epitope and partial overlap with the class 1 epitope proximal to the ACE2 receptor binding site (RBS) but distal to the major protruding ridge of the RBD (**Figures 4A-B**, **Table S2**). This class 1/4 epitope has alternatively been referred as the RBS-D/CR3022 site,^55^ the F2/F3 sites,^56^ or the RBD-6 site,^57^ and it is targeted by noteworthy broad-and-potent human antibodies including DH1047,^58^ ADG20,^55^ S2X259,^59^ and SA55-1239.^56^ Hence, antibodies such as SC27 leverage the characteristics of two sites on the RBD: one targets the RBS region, enabling direct competition with the ACE2 receptor binding, ensuring high potency, while the other focuses on the highly conserved CR3022 cryptic site, providing breadth. Unlike other class 1/4 antibodies, which were obtained from the B cells of individuals who had previously recovered from SARS-CoV 2003 infection, SC27 originated from a SARS-CoV-2 vaccine-elicited IgG lineage.

**Figure 4.**
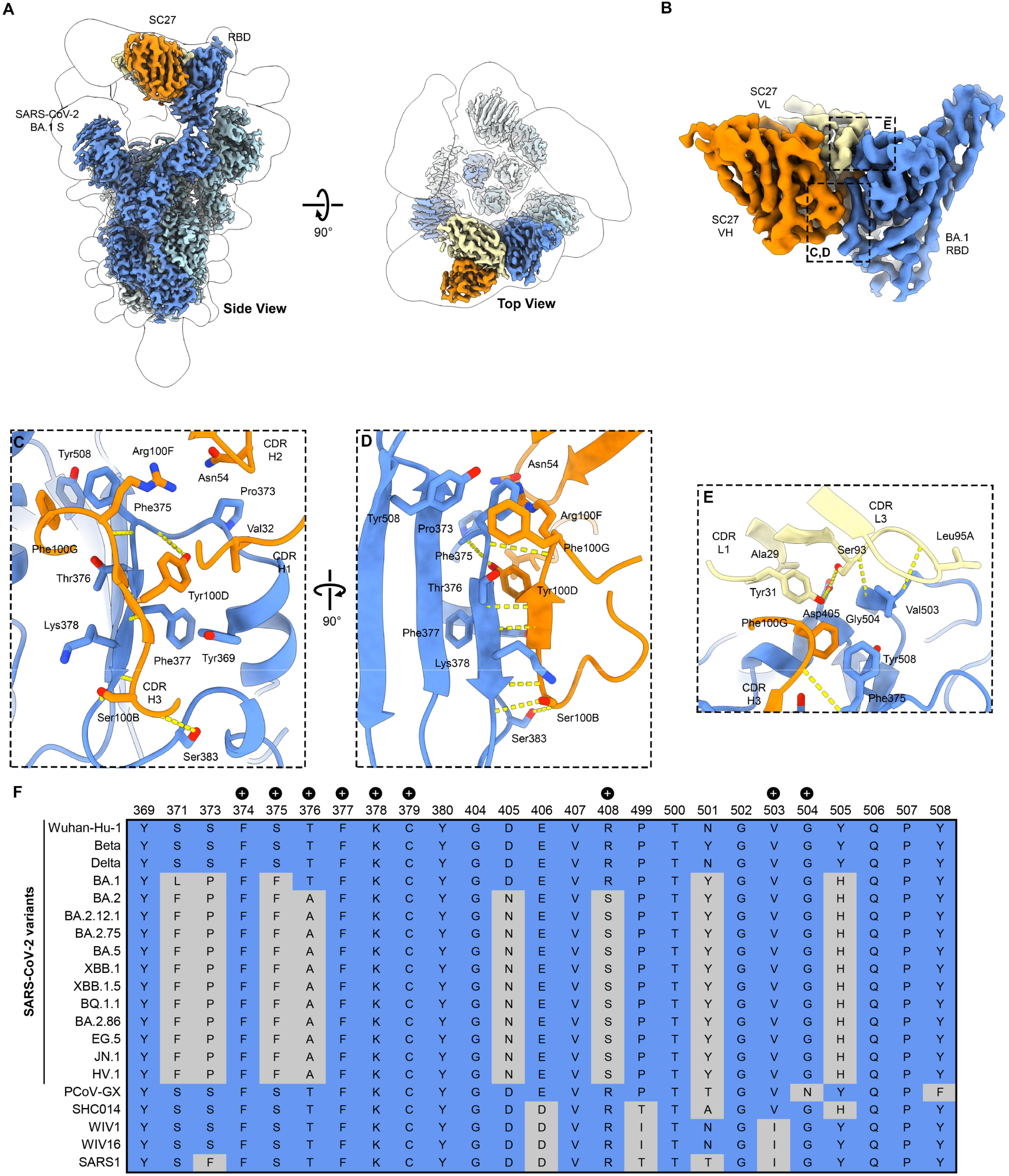
Structural basis for SC27-mediated neutralization. (A) Coulomb potential map of SC27 Fab (orange, yellow) bound to the SARS-CoV-2 BA.1 S protein (blue). (B) Coulomb potential map of SC27 Fab bound to the BA.1 RBD, with boxes encompassing interactions by the heavy chain VH (orange) and light chain VL (yellow) with the RBD. (C-D) Zoomed in view of the interaction between the VH of SC27 (orange) and RBD class 4 epitopic region (blue). (E) Zoomed in view of the interaction of the VL (yellow) of SC27 and RBD class 1 epitopic region (blue). Blue, nitrogen atoms; red, oxygen atoms; dashed yellow lines, hydrogen bonds. (F) Sequence conservation across the RBD– SC27 binding interface (class 1/4 epitope) across an array of SARS-CoV-2 VOCs and zoonotic sarbecoviruses. “+” symbols indicate residues contacted by SC27 via hydrogen bonds.

The binding interface of other class 1/4 mAbs use their antibody VH domain exclusively (or nearly so, except for S2X259) whereas antibody SC27 uses both its VH and its VL domains to make extensive, bispecific contacts. The packing interface of SC27 to its class 1/4 interface buries 717 Å^2^ of surface area on the RBD (542 Å^2^ by the heavy chain and 175 Å^2^ by the light chain) (**Figure 4C-D**). Two residues, Val32 (CDR-H1) and Asn54 (CDR-H2), form van der Waals contacts with Pro373 on the RBD. Notably, the SC27 CDR-H3 adopts a β-strand conformation that forms an antiparallel β-sheet with the spike β2 β-strand composed of residues 375-380, with all possible backbone hydrogen bonds formed over this residue range and two residues that flank it (**Figure 4D**). The intermolecular β-sheet is capped on both ends by additional contacts: at one end, Ser100B in the CDR-H3 forms a sidechain-to-backbone hydrogen bond with Cys379 on the RBD. At the other end, Phe100G and Arg100F form ϖ-ϖ and ϖ-cation interactions, respectively, with RBD Phe375. The β-sheet is further stabilized by a hydrogen bond between the sidechain of Tyr100D (CDR-H3) and the backbone of Phe375 on the spike. The light chain (**Figure 4E**) forms a separate, extensive contact proximal to the class 1 epitope, making four hydrogen bonds with the RBD, two of which are formed between the backbones of the CDR-L3 and Val503 and Gly504 on the RBD. The two other light chain hydrogen bonds (Ser93 and Tyr31) both contact Asp405 on the RBD.

Interestingly, several residues within the SC27 spike epitope have been mutated between the Wuhan-Hu-1 and BA.1 strains, suggesting that SC27 may possess some degree of structural plasticity since it is able to neutralize both with high potency. The potential for SC27 structural flexibility may be accommodated, in part, by the cooperative “bispecific” involvement of both its VH and its VL domains binding to two distinct epitopic regions that are both highly conserved across diverse SARS-CoV-2 variants, SARS-CoV 2003, and bat and pangolin CoVs (**Figure 4F**). This is especially true for conservation of G504, which has been preserved in SARS-CoV-2 ancestral virus and all VOCs, and is constant among CoVs in clades 1a, 1b and 3 (except for Pangolin-GX-2017, which contains N504 instead, although SC27 still recognizes it).

Notably, most epitope residues with a large buried surface area (BSA) when bound by SC27 are highly conserved (**Table S2**). Additionally, key epitope residues with side chains that form hydrogen bonds with SC27 show minimal natural variation **(Figure 4F; Table S2)**. The findings from a high-throughput DMS platform^60^ using replicative pseudotyped lentiviruses to evaluate the effects of S mutations on antibody neutralization and S-mediated infection quantified the probability of XBB.1.5 S mutations to escape SC27 (**Figure S7A-C**) but such mutations (e.g., G504) are rarely found in circulating viruses. Lastly, it is noteworthy that individuals with hybrid immunity, who were donors during the SARS-CoV 2003 outbreak and who subsequently received vaccination with the ancestral S protein of SARS-CoV-2, exhibit an increased prevalence of polyclonal plasma antibodies that share a class 1/4 binding footprint similar to SC27 (**Figure S7D**).

## DISCUSSION

The continuing emergence of globally circulating SARS-CoV-2 variants, coupled with abundant zoonotic reservoirs of pre-emergent sarbecoviruses, highlights the need for a deep understanding of the durability of polyclonal antibody responses (humoral immunological memory) to both viral infections and repeated vaccinations. Based on our data, we conclude that hybrid immunity and serological IgG recall are imprinted and fixed by one’s initial mode of exposure to S protein—and this imprinting differs between natural and vaccine-induced immunity. Immunological imprinting appears to be fundamental in establishing and perpetuating a persistent antibody orientation within humoral immunological memory. Infection shows a pronounced augmentation of S2-subunit-specific IgG antibody lineages, a response that is notably diminished with vaccination alone, consistent with bulk-level serological titers as reported by others.^28,61^

We note, however, that the prevailing evidence in the existing literature does not support the concept of *epidemiological imprinting* at a population level, so our data should not be extrapolated regarding the possibility of differential protection from COVID-19 disease. Clinical data suggest that any prior exposure to SARS-CoV-2, whether through infection or vaccination, significantly reduces the severity of subsequent infections.^62^ Despite the literature not supporting the concept of epidemiological imprinting, the SARS-CoV-2 pandemic represents a paradigm shift where both natural infections coupled with vaccinations were used in parallel to obtain global immunity in a short window. It is reasonable to estimate that the global population has been vaccinated with ∼4 billion killed vaccines and about 2-3 billion mRNA vaccines. Differential targeting may well elicit unique imprinting responses that influence (1) virus evolution rates, (2) durability of the response, and (3) BT potential and mechanism. More studies are warranted, given the potential for a global health pattern to unfold when, in the future, our species may be exposed to another highly transmissible emerging virus.

Yet, there is a clear consensus that current bivalent vaccine boosters, containing either BA.1 or BA.5 Omicron spike sequences, when compared to monovalent vaccine boosters, do not result in significantly higher binding titers or virus-neutralizing titers against the SARS-CoV-2 variants.^63–66^ Our research findings based on IgG molecular serology are consistent with data from analyses of the binding and neutralization breadth of antibodies cloned from S-specific memory B cells^67,68^ and strongly indicate that the presence of the ancestral spike, whether within existing COVID-19 vaccine formulations or introduced through an early-wave pandemic infection, is a key factor in inducing so-called *deep immunological imprinting* or *hybrid immune damping*.^17,63^

Extensive efforts have led to the identification and structural characterization of dozens of broadly neutralizing antibodies (bNAbs) to SARS-CoV-2, the majority of which have now been escaped by evolved variants such as Omicron BA.1. Overwhelmingly, these bNAbs were identified by the cloning of immunoglobulin variable (IgV) region genes from peripheral blood memory B cells. Here, we have reported the first cloning of a SARS-CoV-2 bNAb isolated from polyclonal IgG at the level of the secreted (plasma) proteome. The progenitor of bNAb SC27 was first recognized as a circulating IgG induced by the Pfizer/BioNTech mRNA vaccine in a naïve donor. SC27 experienced affinity maturation, demonstrated by a notable rise in somatic IgHV-gene mutations over time, and exhibited heightened abundance due to an ensuing BT infection. SC27 showcases exquisitely high binding affinity (<5 pM intrinsic affinity as a monovalent F_ab_) validated against both ancestral and BA.1 strains and demonstrates strong neutralizing potency against SARS-CoV 2003, clade 1a and 1b zoonotic bat and pangolin viruses, and against all major clade 1b SARS-CoV-2 Omicron variants, including the most recent Omicron VOC, BA.2.86. Lastly, SC27 provides *in vivo* protection against viral challenge in the MA10 mouse model. This case offers clear molecular-level evidence for how hybrid immunity can elevate serological function and enhance memory through the repeated elicitation and ongoing sculpting of a single IgG plasma antibody.

Antibody SC27 also marks the first demonstration of a class 1/4 antibody present as a circulating IgG (and not found solely within the B-cell compartment). It has been proposed that vaccines and mAbs that focus on the comparatively stable class 1/4 neutralizing antibody epitope may offer potential protection against future zoonotic transmission events and emerging variants of SARS-CoV-2.^69^ The fact that other class 1/4 mAbs were all derived from SARS-CoV 2003 convalescent donors was presumed to explain why BCR-derived mAbs directed to this site are seemingly so rare. SC27, however, was generated by mRNA vaccination of a naïve donor who had no detectable titers against either the SARS-CoV-2 S or nucleocapsid (N) proteins. Importantly, SC27’s functional potency and its abundance were both augmented in this donor by a hybrid immunological event (BT infection). In this regard it is interesting to note that polyclonal plasma antibodies, sharing the SC27-like class 1/4 binding footprint, become enriched in hybrid immune SARS-CoV 2003 donors when they receive vaccination by SARS-CoV-2 ancestral S protein.

A four-to-five amino acid stretch in the SC27 CDR-H3 adopts a β-strand conformation, making backbone contacts with the RBD β2 strand to extend the internal β-sheet located on the cryptic underside of the receptor. The β2 strand is a linear peptide spanning S residues 369 to 386 and is involved in all class ¼ binding interactions. The close juxtaposition and the complementariness^70^ of surface topology between the antibody CDR-H3 and the RBD β2 strand suggests the possibility that the intrinsic geometry of the β-sheet conformation helps facilitate the broad binding and neutralization capabilities of SC27. Side chains on all β-strands point perpendicular to the hydrogen bonds that form between strands.^71^ In this manner, potential escape mutations might have limited capacity to directly interfere with the SC27 binding interface. Deep mutational scanning indicated the potential for SARS-CoV-2 spike mutations to evade SC27, yet such genetic alterations are seldom identified in viruses circulating in the population and the conservation of G504 is conserved across 3 of the 4 sarbecovirus clades (1a, 1b and 3). Despite the accumulation of mutations in the RBD among VOCs, the secondary and tertiary structure of the spike RBD has been conserved since SARS-CoV-2 emerged in 2019. For example, the Wuhan-Hu-1 (PDB 6VSB) and XBB.1.5 (PDB 8SPI) spikes share a high degree of structural similarity, with an RMSD equal to 0.73 Å over all Cα atoms in the RBD. Antibodies that exhibit β-sheet–like binding properties, as well as other backbone-backbone contacts such as those in the SC27 CDR-L3, might thus be uniquely situated to adapt to emerging SARS-CoV-2 variants. Indeed, the β-strand-like binding of SC27 has been observed for broadly neutralizing antibodies directed at other β-CoVs^72^ and HIV^73^.

Based upon our observations of IgG anti-S repertoires in vaccinated BT donors, our results suggest that deploying diverse RBDs in next-generation vaccine strategies (to emulate hybrid immunological S scenarios) should elicit SC27-like class 1/4 antibodies that have lengthy CDR-H3 loops juxtaposed with the RBD β-sheet backbone to confer broad reactivity while the orientation of their light chains should contribute to steric blockade of the ACE2 binding site. Hence, vaccines capable of triggering these antibodies^69,74,75^ might offer protection against SARS-CoV-2, its variants, and newly emerging zoonotic sarbecoviruses, eliminating the necessity for frequent updates in response to new outbreaks.

## LIMITATIONS OF THE STUDY

Limitations include the following: (i) small number of individuals which may limit the interpretation of the data and leaves it unclear how common the SC27-like antibody might be in human populations; (ii) IgG subclasses were not examined, which precludes accurate future study of Fc-mediated effector functions, like ADCC or ADCP, rather than merely ascertain antibody neutralization activity as reported here; and (iii) immunological imprinting of mucosal immune responses were not examined.

## Acknowledgments

The authors thank D. Billick for his assistance in drawing blood used in this project. This research was chiefly funded by a National Institutes of Health (NIH) COVID-19 SeroNet grant awarded to R.S.B. with subaward to G.C.I. (U54-CA260543). Major additional funding was awarded by the NIH to J.D.B. (R01-AI141707), J.S.M. (R01-AI127521), and by the Bill and Melinda Gates Foundation grants INV-004956 (G.G. and J.J.L.) and INV-031624L (J.S.M., R.S.B. and G.C.I.). J.D.B. is an Investigator of the Howard Hughes Medical Institute. We are grateful to the Biological Mass Spectrometry Facility and the Genome Sequencing & Analysis Facility at the University of Texas at Austin.

## Author contributions

Conceptualization: W.N.V., G.G., R.S.B., J.J.L. and G.C.I.; Methodology: W.N.V., M.A.M., P.O.B., J.M.M., J.M.P., S.R.L., B.D., P.L., J.J.L. and G.C.I.; Investigation: W.N.V., M.A.M., P.O.B., J.M.M., S.A.K., J.M.P., S.R.L., B.D., D.R.T., J.K., Y.H., E.S., I.N.C., M.M., C.P., J.E.M., T.S., A.S., P.L., and J.J.L.; Data Analysis and Interpretation: W.N.V., M.A.M., P.O.B., J.J.M., S.A.K., J.M.P., S.R.L., B.D., P.L., J.D.B., J.J.L. and G.C.I.; Data Curation: W.N.V., M.A.M. and P.O.B.; Writing: Original Draft, W.N.V., P.O.B., J.J.L. and G.C.I.; Writing: Review & Editing: W.N.V., M.A.M., G.G., R.S.B., J.J.L. and G.C.I.; Resources: P.L., J.D.B., G.G., J.S.M., R.S.B., J.J.L. and G.C.I.; Funding: J.D.B., G.G., J.S.M., R.S.B., J.J.L. and G.C.I.

## Competing interests

R.S.B. is a member of advisory boards for VaxArt, Takeda and Invivyd, and has collaborative projects with Gilead, J&J, and Hillevax, focused on unrelated projects. S.R.L. and R.S.B. are co-inventors of methods and uses of mouse-adapted SARS-CoV-2 viruses (US patent US11225508B1). J.D.B. is on the scientific advisory boards of Apriori Bio, Invivyd, Aerium Therapeutics and the Vaccine Company. J.D.B. and B.D. are inventors on Fred Hutchinson Cancer Center licensed patents related to viral deep mutational scanning. R.S.B., G.G., J.J.L., G.C.I., and W.N.V. are inventors on a provisional U.S. patent application for mAb SC27 and other new antibodies described in this manuscript, entitled ‘‘Broadly neutralizing human monoclonal antibodies that target the SARS-CoV-2 receptor binding domain (RBD)’’ (63/491,270).

---

**Figure S1.**
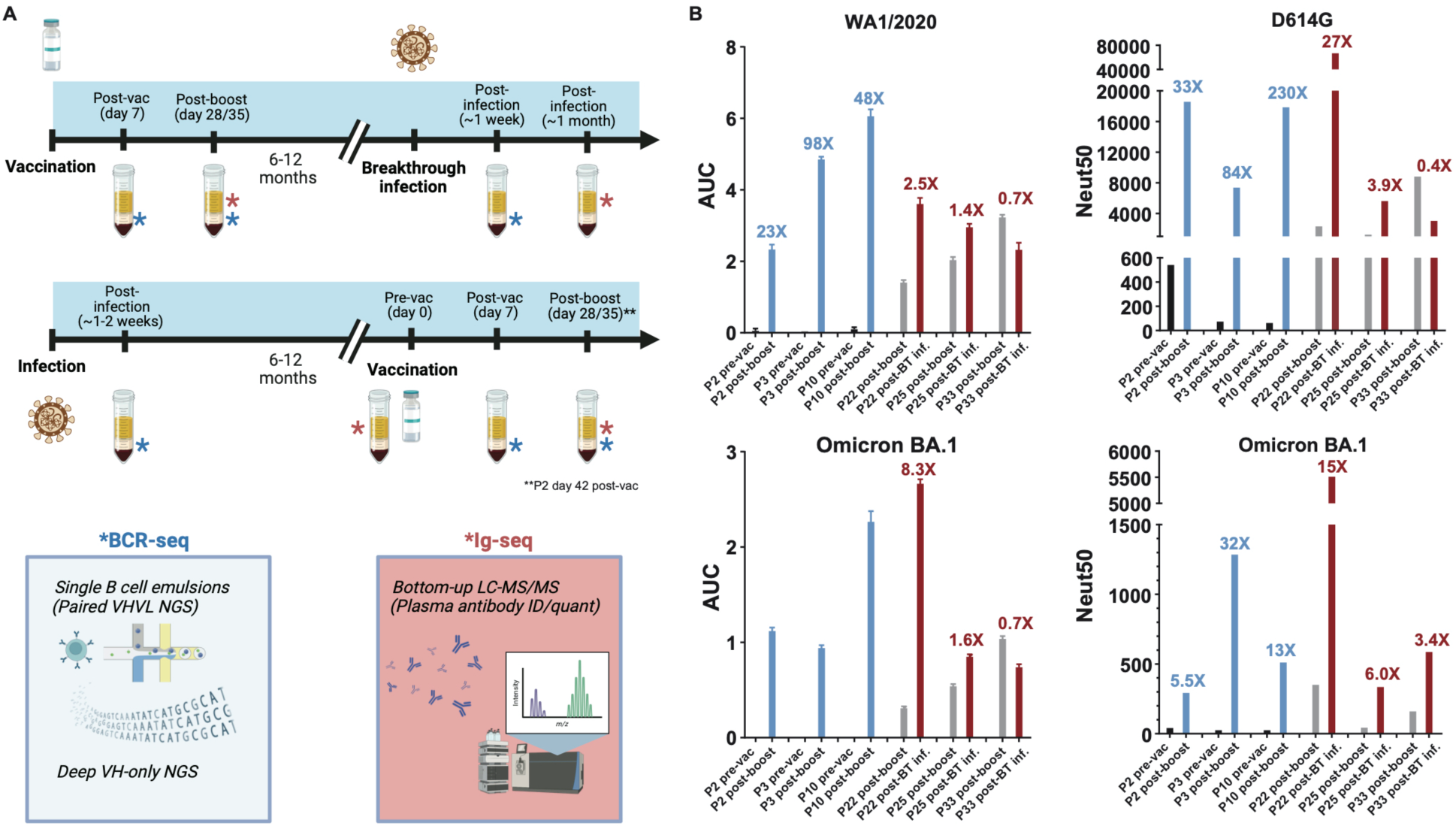
Experimental workflow and bulk serology of a select cohort. (**A**) Sample and workflow schematic for vaccination-infection and infection-vaccination groups examined in this study. (B) Bulk serological spike binding (left) and viral neutralization (right) titers against ancestral (top) and Omicron BA.1 (bottom) SARS-CoV-2 viruses across six donors at each Ig-seq time point examined.

**Figure S2.**
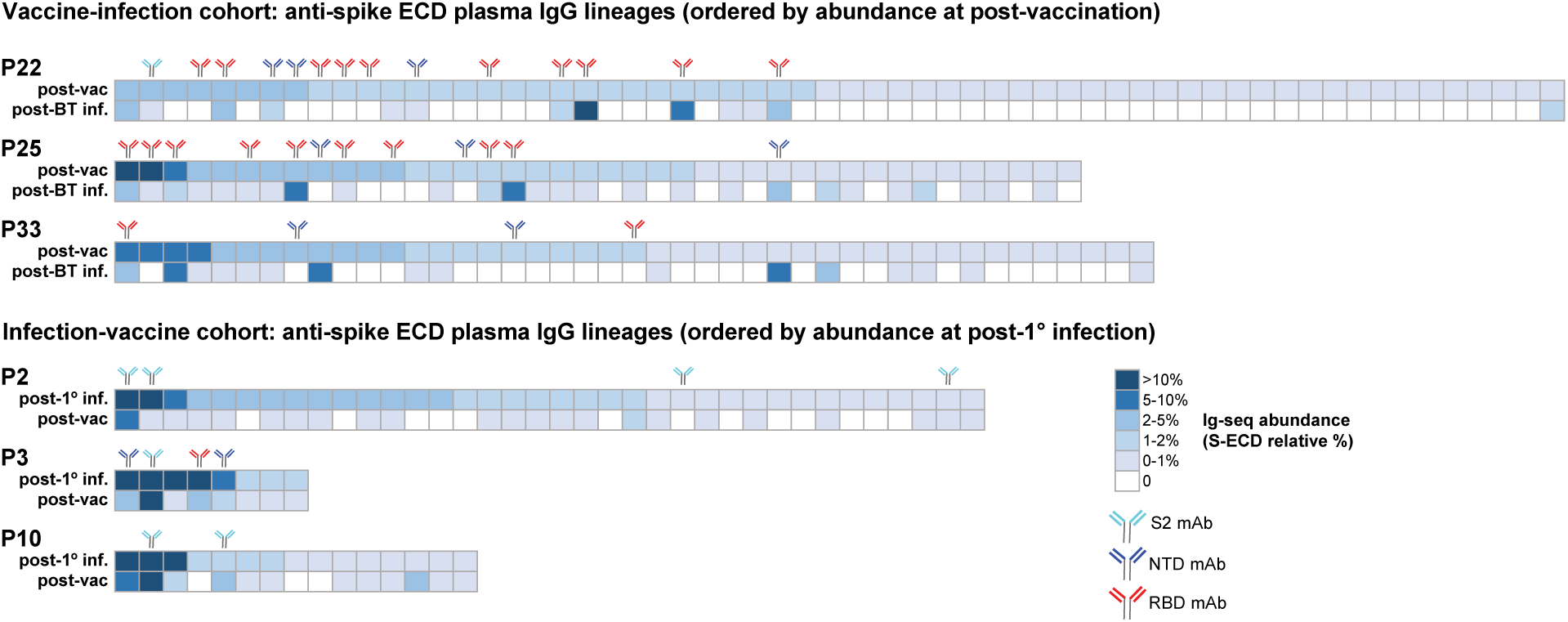
Ig-Seq heatmap of IgG anti-S hybrid immunity. For each of the six donors examined by Ig-Seq, each column represents a unique plasma IgG lineage across the two time points examined, ordered by relative abundance (% anti-spike plasma IgG) at the initial time point. Colored antibody markers above each lineage indicate full length VH:VL pairs were identified, and recombinant antibody was expressed, with domain specificity determined by indirect ELISA against individual spike domains. Only plasma IgG lineages present at >0.5% relative abundance at the initial time point are shown.

**Figure S3.**
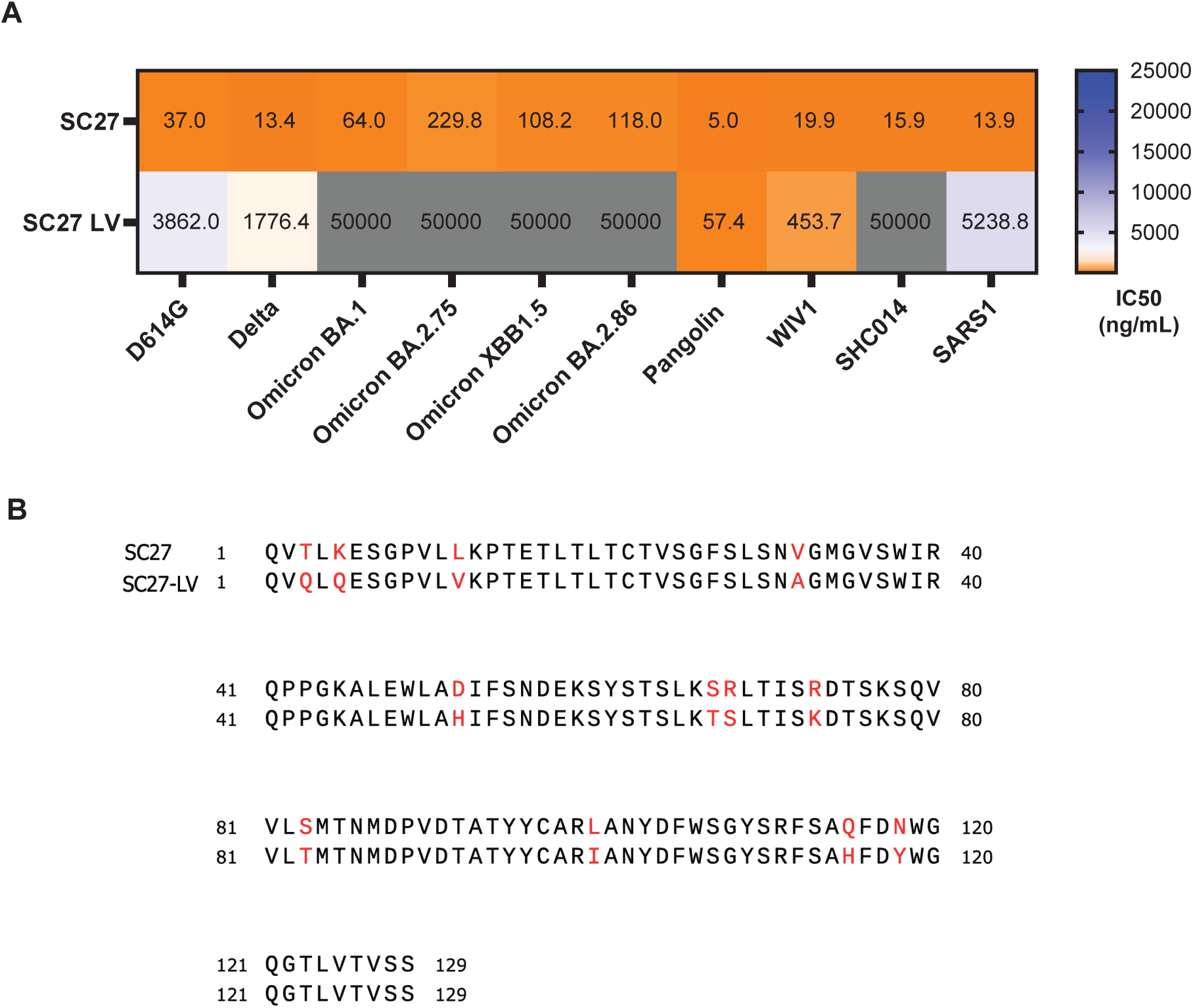
Comparison of SC27-LV mAb clone, identified in plasma post-vaccination, to SC27 (post-BT infection clone). (A) Live-virus neutralization assays using SC27-LV and SC27 screened against a panel of SARS-CoV-2 VOCs as well as zoonotic sarbecoviruses. (B) Sequence alignment of SC27 and SC27-LV, with amino acid differences highlighted in red. “LV” notates “late vaccination” based upon level of somatic hypermutation within the SC27 plasma IgG lineage.

**Figure S4.**
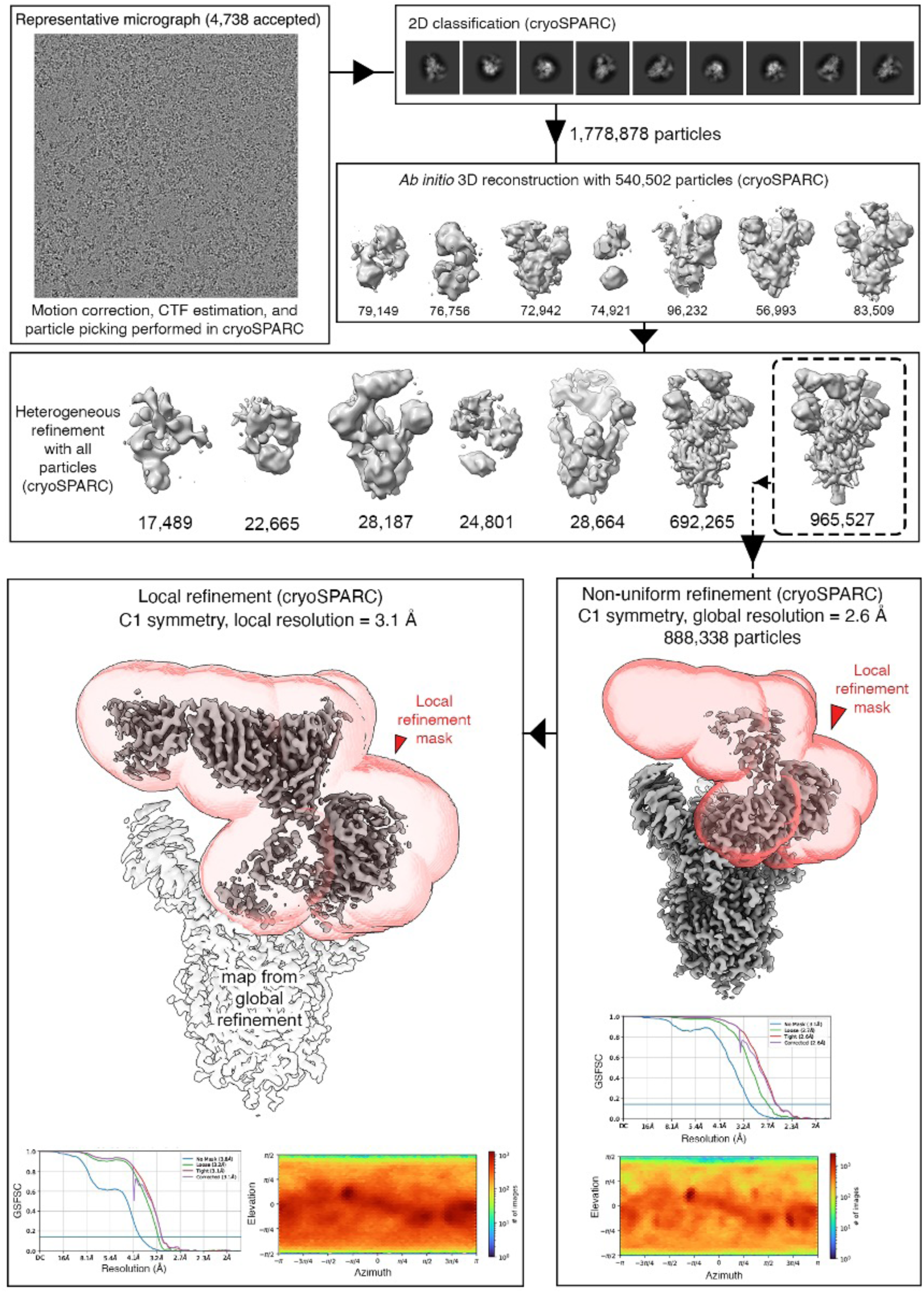
Cryo-EM processing summary for the spike trimer bound to SC27 Fab.

**Figure S5.**
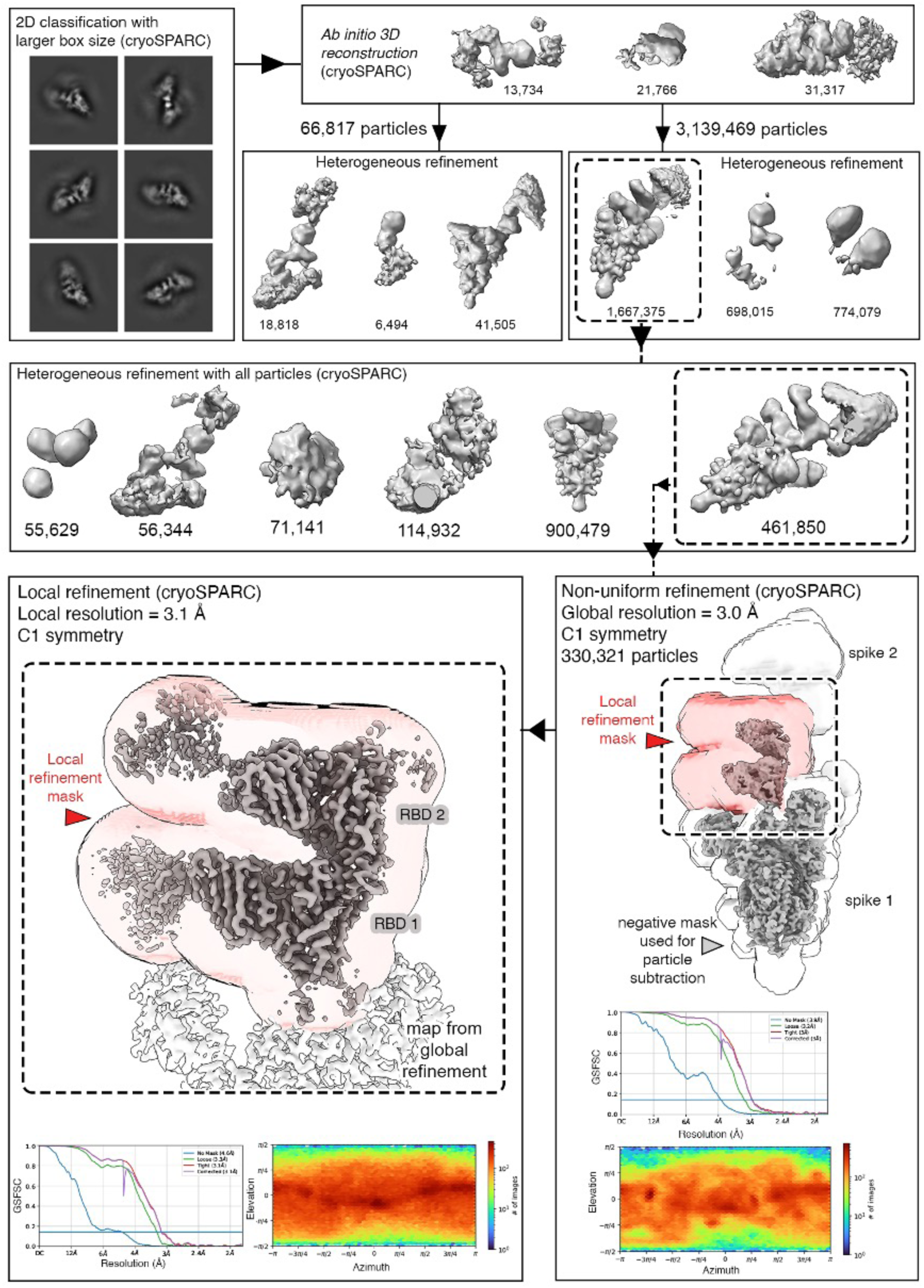
Cryo-EM processing summary for the RBD-mediated dimer-of-trimers.

**Figure S6.**
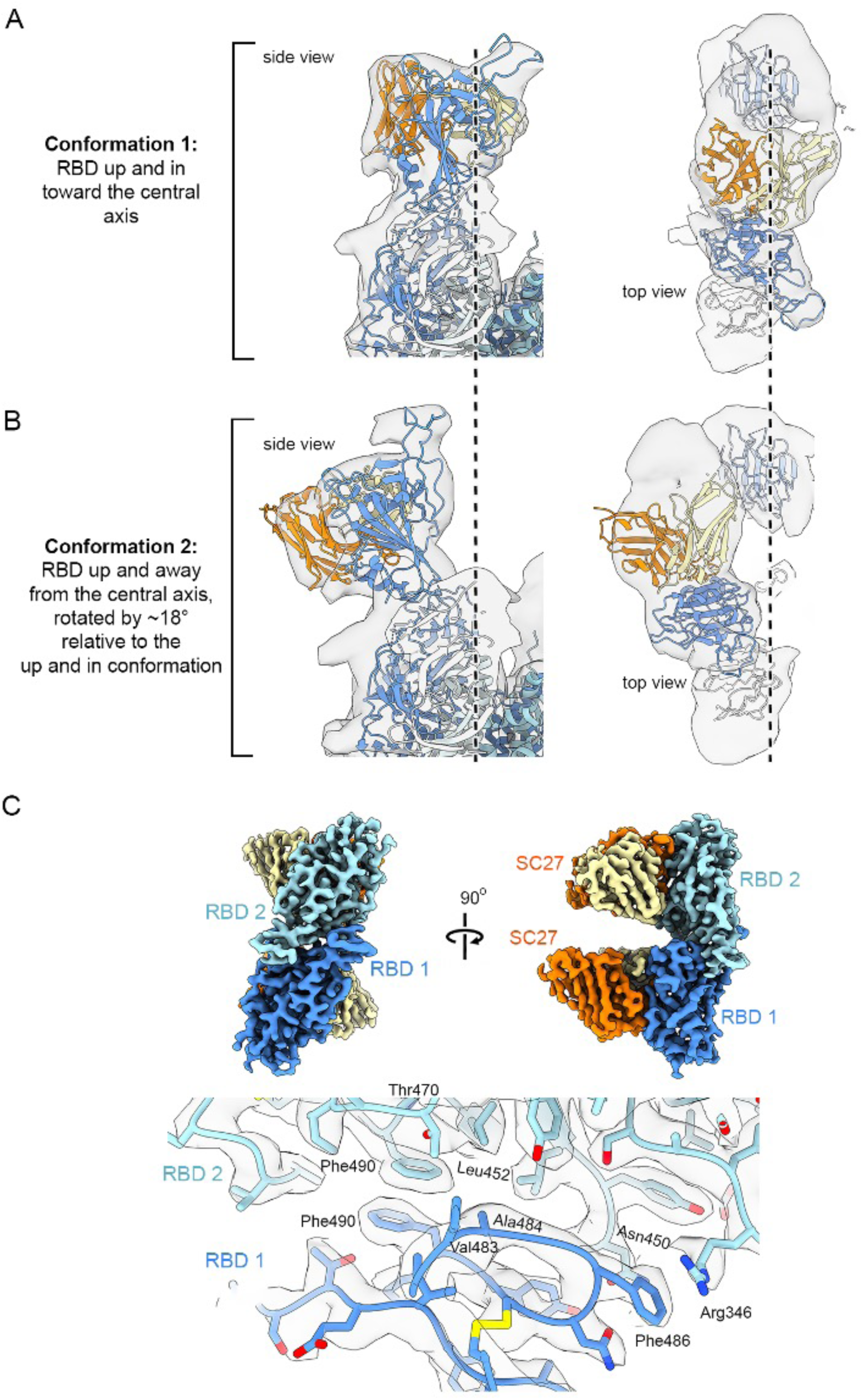
Additional Cryo-EM processing reveals RBD conformational heterogeneity.

**Figure S7.**
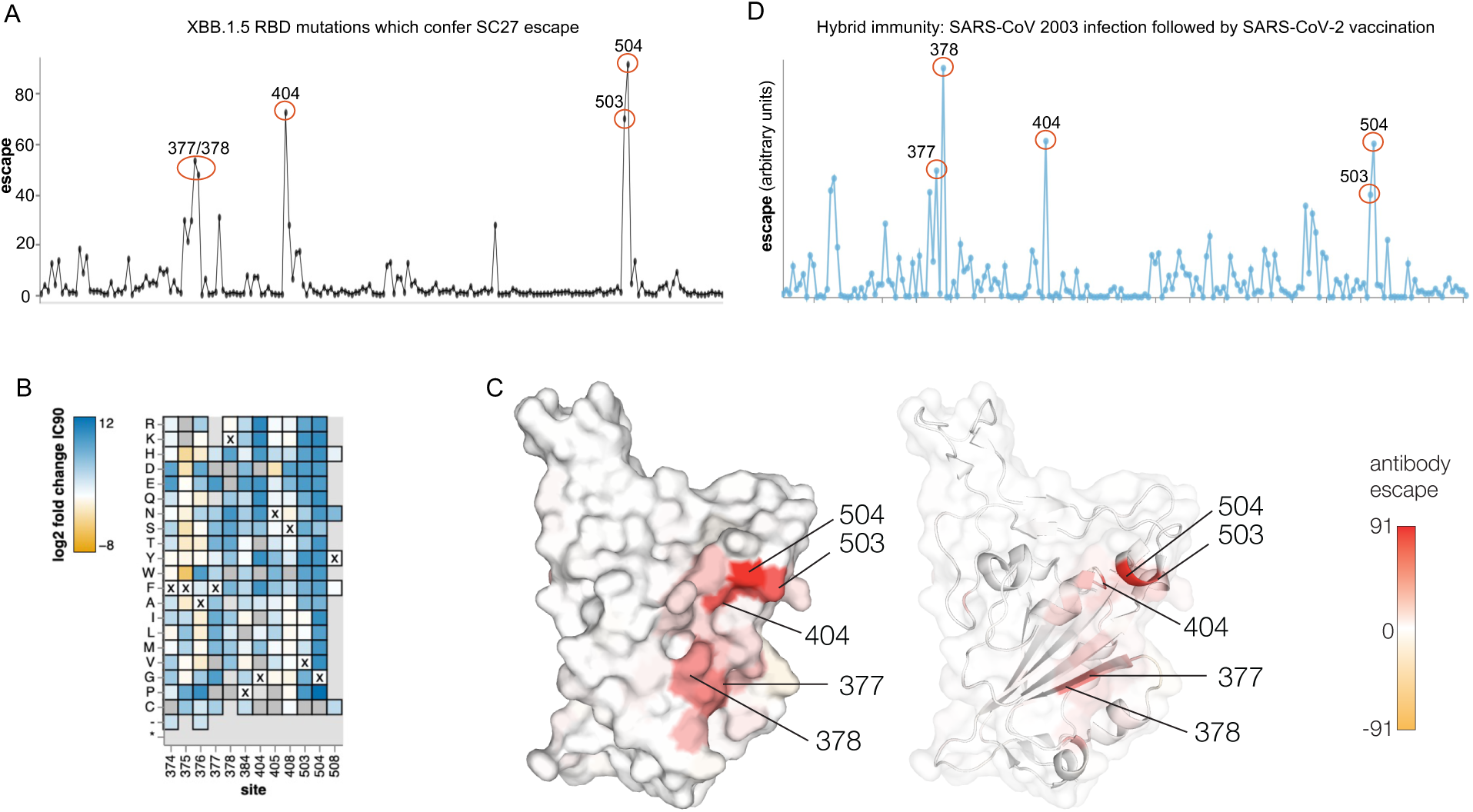
Deep mutational scanning of SARS-CoV-2 XBB.1.5 RBD and SC27 mAb escape. (A) Total escape scores at each site in the XBB.1.5 RBD. (B) Heatmap of mutation-escape scores at key sites. Residues marked with X are the wild-type amino acids in XBB.1.5. Amino acids absent from the library are shown in grey. Complete line plot, heatmaps and code for (A) and (B) can be found at https://dms-vep.github.io/SARS-CoV-2_XBB.1.5_RBD_DMS_SC27/htmls/summary_overlaid.html. (C) Surface representation of spike RBD colored by the sum of escape scores at that site. (D) Key sites targeted in hybrid immunity by human antibodies elicited by SARS-CoV 2003 infection followed by SARS-CoV-2 vaccination (Aggregated data available in the Bloom Lab antibody escape calculator here: https://jbloomlab.github.io/SARS2-RBD-escape-calc/).

**Table S1.** Convalescent and post-vaccination donor blood sample information. Lowercase “i” following donor name indicates primary infection prior to first vaccination, “v” indicates vaccination after primary infection, “nv” indicates naïve vaccination, and “BT” indicates breakthrough infection after vaccination. For vaccines administered in two doses (Pfizer and Moderna), “INF/VAX date” refers to date of first infection or first vaccine dose. “Days since previous S exposure” refers to the number of days between infection and vaccination, in either order. In the “Blood draw time point/s” column, “D# post-vax” refers to the number of days after the first vaccine dose. “PSO” = “post-symptom onset”.

| Donor | Sample | Type | Vaccine | INF/VAX date | Days since previous S exposure | Blood draw time point/s |
| --- | --- | --- | --- | --- | --- | --- |
| P2 | P2i | Primary infection | N/A | Mar 10, 2020 | N/A | ~1-2 weeks PSO |
| P3 | P3i | Primary infection | N/A | Mar 9, 2020 | N/A | ~1-2 weeks PSO |
| P10 | P10i | Primary infection | N/A | Jul 15, 2020 | N/A | ~1-2 weeks PSO |
| P2 | P2v | Post-infection vaccination | Janssen JNJ 78436735 | Apr. 9, 2021 | 395 | D0, D7, D42 post-vax |
| P3 | P3v | Post-infection vaccination | Pfizer BNT162b2 | Mar. 24, 2021 | 380 | D0, D7, D28/35 post-vax |
| P10 | P10v | Post-infection vaccination | Pfizer BNT162b2 | Apr. 7, 2021 | 266 | D0, D7, D28/35 post-vax |
| P22 | P22nv | Naïve vaccination | Pfizer BNT162b2 | Jan. 27, 2021 | N/A | D7, D28/35 post-vax |
| P25 | P25nv | Naïve vaccination | Pfizer BNT162b2 | Feb. 24, 2021 | N/A | D7, D28/35 post-vax |
| P33 | P33nv | Naïve vaccination | Moderna mRNA 1273 | Feb. 10, 2021 | N/A | D7, D28/35 post-vax |
| P22 | P22BT | Breakthrough infection | N/A | Jul. 5, 2021 | 159 | ~1 week, ~1 month PSO |
| P25 | P25BT | Breakthrough infection | N/A | Jul. 27, 2021 | 153 | ~1 week, ~1 month PSO |
| P33 | P33BT | Breakthrough infection | N/A | Oct. 11, 2021 | 243 | ~1 week, ~1 month PSO |

**Table S2.**
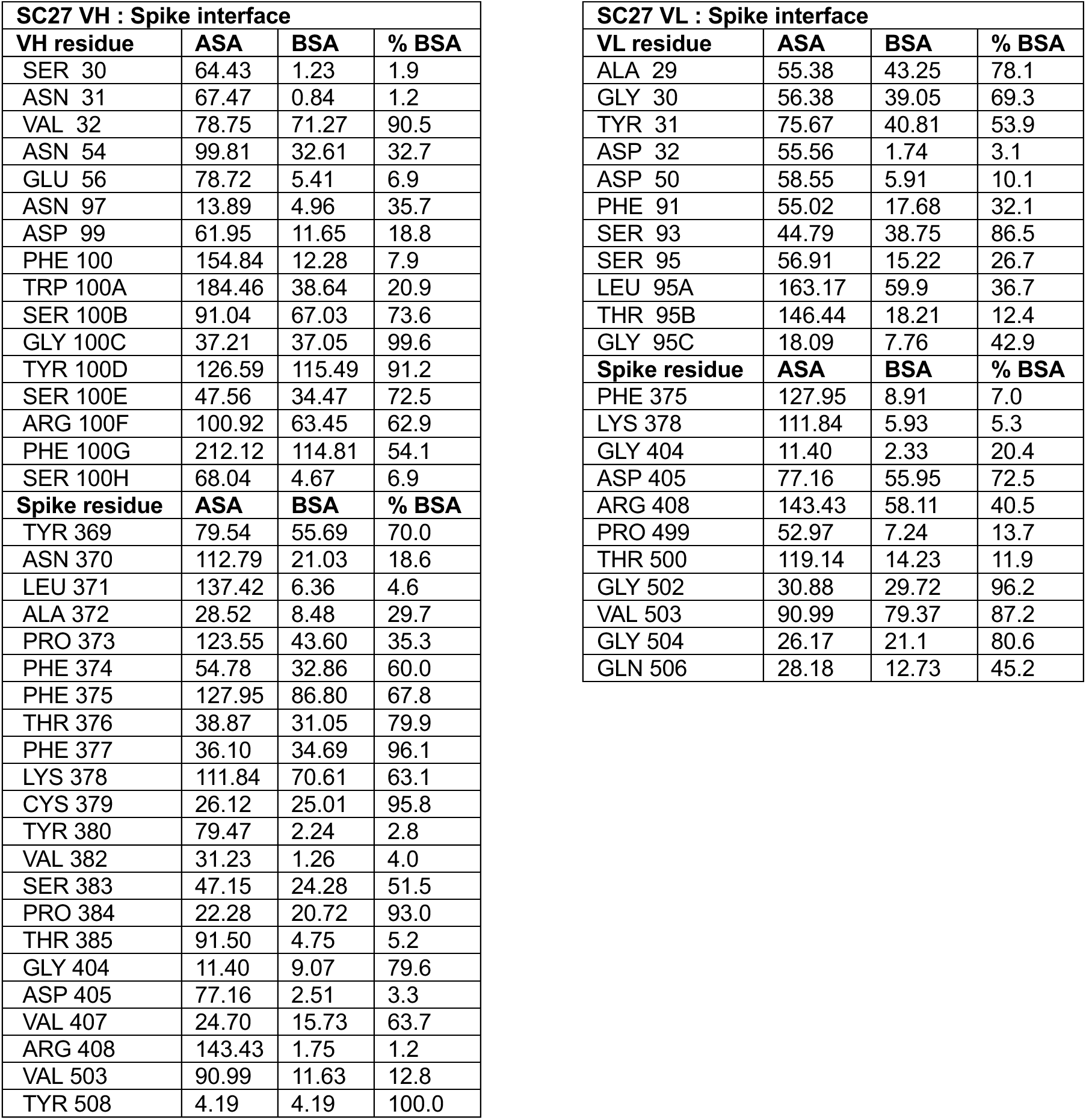
PISA analysis of interface between SC27 F_ab_ and Omicron BA.1 spike protein. “ASA” = “accessible surface area”. “BSA” = “buried surface area”. “% BSA” = percentage of buried surface area for each residue within the interface. Only residues with interface BSA >0 are listed. All surface area values are Å^2^.

**Table S3.**
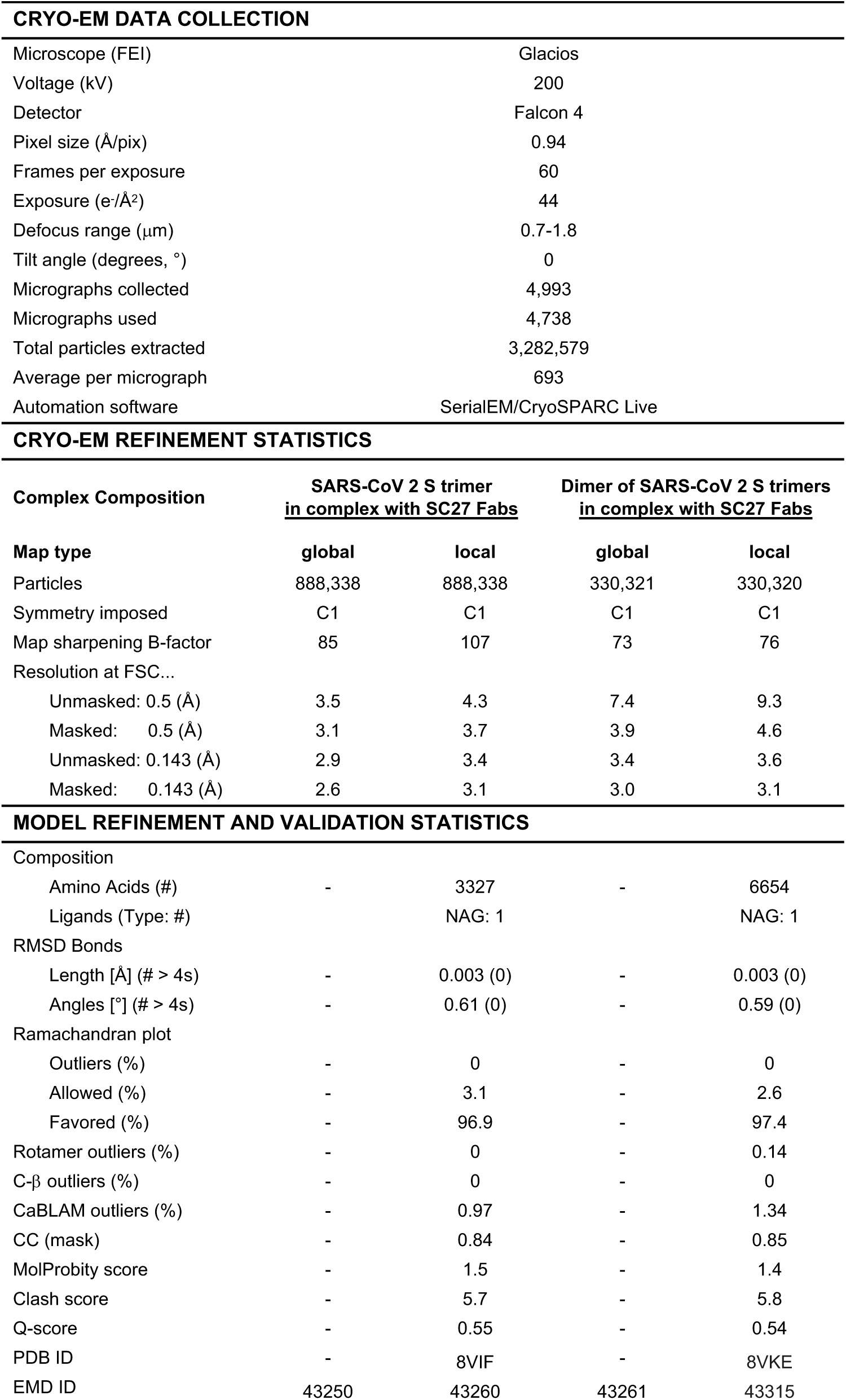
Cryo-EM data collection and refinement statistics.

## Materials and Methods

### Donor cohort and collection of blood plasma and PBMCs

The SARS-CoV-2 immune plasmas used in this study were collected from i) convalescent non-hospitalized PCR-confirmed individuals with symptomatic disease (P2i, P3i, P10i), ii) SARS-2 naïve individuals vaccinated with either the Pfizer BNT162b2 or Moderna mRNA-1273 SARS-2 vaccines (P22, P25, P33), iii) previously infected individuals vaccinated with either the Pfizer BNT162b2 or Janssen JNJ-78436735 SARS-2 vaccines, or iv) previously vaccinated, but SARS-2 infection naïve individuals that experienced a SARS-CoV-2 breakthrough infection. Whole blood was separated into a plasma and PBMC fractions via density gradient centrifugation using Histopaque-1077 media (Sigma-Aldrich).

### Expression and purification of SARS-CoV-2 proteins

For Ig-Seq antibody proteomics, the cloning, expression, and purification of the prefusion-stabilized spike ectodomain HexaPro spike variant^1^ containing substitutions S383C and D985C^2^ with a C-terminal TwinStrep tag, have been previously described.^3,4^

For indirect ELISAs to identify mAbs specific to the SARS-CoV-2 spike S2 domain, a stabilized construct of the S2 subunit (S2-37, a kind gift from Jason McLellan) was expressed by transiently transfecting transiently transfecting plasmid using FreeStyle 293-F cells (Thermo Fisher) using polyethyleneimine, with 5 µM kifunensine being added 3h post-transfection. The cell culture was harvested four days after transfection and the medium was separated from the cells by centrifugation. Supernatants were passed through a 0.22 µm filter followed by passage over StrepTactin resin (IBA). The sample was further purified by size-exclusion chromatography using a Superose 6 10/300 column (GE Healthcare) in PBS.

For mAb RBD competition ELISAs to determine RBD class specificity of mAbs isolated from donors, recombinant RBD-spike domain 1 (RBD-SD1) tagged with human IgG Fc was transiently transfected using FreeStyle 293-F cells and purified using Protein G Plus Agarose (Pierce Thermo Fisher Scientific). RBD-SD1 was cleaved from the Fc by incubation with 3C protease (produced in-house) overnight at 4 °C. 3C protease was removed using Ni-NTA resin and RBD-SD1 was buffer-exchanged into PBS and concentrated using 10,000 MWCO Vivaspin centrifugal spin columns (Sartorius).

For ACE2 inhibition assays and cryo-EM structural analysis, two variants of SARS-CoV-2 spike, Wuhan-Hu-1 and BA.1, were produced using plasmids encoding the spike ectodomain with a C-terminal T4 fibritin foldon trimerization motif, 8X His tag, and TwinStrep tag. Briefly, plasmids were transiently transfected into HEK293F cells using polyethyleneimine (25,000 Da) at a mass ratio of 9:1 (PEI:DNA). Cells were grown in suspension for 6 days shaking at 37 °C, 8% CO_2_. The media was harvested by centrifugation, then concentrated by tangential flow filtration while buffer exchanging into 1X PBS. The concentrated medium was passed over Strep-Tactin Superflow Resin (IBA Life Sciences), washed with 1X PBS, and eluted with the manufacturer’s Buffer E, which contains desthiobiotin. Elution fractions were analyzed by SDS-PAGE. Fractions containing spike were pooled, concentrated by centrifugal filtration, and separated by size exclusion chromatography (SEC) (Superose 6 Increase columns) using running buffer containing 2 mM Tris pH 8 and 200 mM NaCl supplemented with 0.01% sodium azide (w/v). Trimeric spike fractions were pooled, concentrated in centrifugal filters, aliquoted, flash frozen in liquid nitrogen, and stored at -80 °C.

### V_H_ repertoire sequencing

PBMCs were lysed in TRIzol Reagent (Invitrogen) and total RNA was extracted using RNeasy (Qiagen). First strand cDNA was synthesized from 500 ng mRNA using SuperScript IV (Invitrogen), and cDNA encoding the V_H_ regions of the IgG, IgA, and IgM repertoires was amplified with a multiplex primer set^5^ using the FastStart High Fidelity PCR System (Roche) under the following conditions: 2 min at 95 °C; 4 cycles of 92 °C for 30s, 50 °C for 30s, 72 °C for 1 min; 4 cycles of 92 °C for 30s, 55 °C for 30s, 72 °C for 1 min; 22 cycles of 92 °C for 30s, 63 °C for 30s, 72 °C for 1 min; 72 °C for 7 min; hold at 4 °C, as previously described.^5^ Products were sequenced by 2x300 paired-end Illumina MiSeq.

### Paired V_H_:V_L_ repertoire sequencing

PBMCs were co-emulsified with oligo d(T)_25_ magnetic beads (New England Biolabs) in lysis buffer (100 mM Tris pH 7.5, 500 mM LiCl, 10 mM EDTA, 1% lithium dodecyl sulfate, and 5 mM dithiothreitol) using a custom flow-focusing device as previously described.^6^ The magnetic beads were washed, resuspended in a one-step RT-PCR solution with an overlap extension VH and VL primer set as previously described,^6^ emulsified using a dispersion tube (IKA), and subjected to overlap-extension RT-PCR under the following conditions: 30 min at 55 °C followed by 2 min at 94 °C; 4 cycles of 94 °C for 30s, 50 °C for 30s, 72 °C for 2 min; 4 cycles of 94 °C for 30s, 55 °C for 30s, 72 °C for 2 min; 32 cycles of 94 °C for 30s, 60 °C for 30s, 72 °C for 2 min; 72 °C for 7 min; hold at 4 °C. Amplicons were extracted from the emulsions, further amplified using a nested PCR as previously described^6^, and sequenced using 2x300 paired-end Illumina MiSeq.

### Ig-Seq sample preparation and mass spectrometry

Total IgG was isolated from 1 mL plasma via affinity chromatography using Protein G Plus Agarose (Pierce Thermo Fisher Scientific) and cleaved into F(ab’)_2_ fragments using IdeS. SARS-CoV-2 spike-specific F(ab’)_2_ was isolated by affinity chromatography using recombinant antigen (1 mg HexaPro) coupled to 0.05 mg dry NHS-activated agarose resin (Thermo Fisher Scientific) as follows. F(ab’)_2_ (10 mg mL^-1^ in PBS) was rotated with antigen-conjugated affinity resin for 1h, loaded into 0.5 mL spin columns, washed 12X with 0.4 mL Dulbecco’s PBS, and eluted with 0.5 mL fractions of 1% formic acid. IgG-containing elution fractions were concentrated to dryness in a speed-vac, resuspended in ddH2O, combined, neutralized with 1 M Tris / 3 M NaOH, and prepared for liquid chromatography–tandem mass spectrometry (LC-MS/MS) as described previously^7^ with the following modifications: (i) peptide separation using acetonitrile gradient was run for 120 min,(ii) data was collected on an Orbitrap Fusion (Thermo Fisher Scientific) operated at 120,000 resolution using HCD (higher-energy collisional dissociation) in topspeed mode with a 3s cycle time, and (iii) 45s dynamic exclusion of precursors after n=2 fragmentation events in 30s window.

### Bioinformatic analysis

Raw Illumina MiSeq output sequences were trimmed according to sequence quality using Trimmomatic^8^ and annotated using MiXCR.^9^ Sequences with ≥ 2 reads were clustered into clonal lineages using single linkage hierarchical clustering, with clonality defined by 90% CDR-H3 amino acid identity. LC-MS/MS search databases were prepared as previously described,^7^ using custom Python scripts (available upon request). MS searches and MS data analyses were performed as previously described,^7^ adjusting the stringency of the elution XIC:flowthrough XIC filter to 2:1.

### Antibody expression and purification

Cognate V_H_ and V_L_ antibody sequences of interest were synthesized and cloned into a customized pcDNA 3.4 vector containing a human IgG1 Fc region by GenScript Biotech. V_H_ and V_L_ plasmids were mixed at a 1:3 ratio and transfected into Expi293F cells (Thermo Fisher Scientific), which were cultured at 37 °C, 8% CO_2_ for 5 days, then centrifuged at 1000 x g for 10 min. Antibodies were isolated from filtered supernatants using Protein G Plus Agarose (Pierce Thermo Fisher Scientific) affinity chromatography, washed with 20 column volumes of PBS, eluted with 100 mM glycine-HCl pH 2.5, and neutralized with 1 M Tris-HCl pH 8.0. The antibodies were buffer-exchanged into PBS and concentrated using 10,000 MWCO Vivaspin centrifugal spin columns (Sartorius).

For structural studies, SC27 Fab was purified by SEC (Superdex 200 Increase columns) as described above. Fractions containing SC27 Fab were pooled, concentrated using centrifugal filters, aliquoted, flash frozen in liquid nitrogen, and stored at -80 °C.

### ELISA

The methods for enzyme-linked immunosorbent assay (ELISA) to measure anti-SARS-CoV-2 IgG plasma antibody titers have been previously described.^10^ For analysis of mAb reactivity towards different SARS-2 variant recombinant spike ECD proteins, and for the determination of domain binding specificity against recombinant spike RBD, NTD, and S2 proteins, a standard indirect ELISA was used. Costar high binding 96-well assay plates (Corning) were coated with antigens (4 µg ml^-1^) in PBS. Antigens included in-house produced ancestral SARS-COV-2 spike ECD (HexaPro), SARS-CoV-2 B.1.1.529 (Omicron BA.1) variant spike ECD (NCI Serological Sciences Network for COVID-19 [SeroNet]), prefusion stabilized S2 subunit (S2-37), and commercially obtained ancestral SARS-CoV-2 RBD (Bio-Techne) and NTD (Sino Biological) subunits. Antigen-reactive mAbs were detected with goat anti-human IgG (Fab)-horseradish peroxidase (Sigma-Aldrich) conjugate diluted in PBS at a ratio of 1:5000. After washing with 0.1% PBST, the bound antibody was detected with 3,3′,5,5′-tetramethylbenzidine soluble substrate (Ultra TMB; Millipore) using a Synergy H1 Microplate Reader (BioTek Instruments, Inc.).

For plasma RBD competition ELISAs, excess RBD (Bio-Techne; 100 µg mL^-1^ final concentration across serial dilution) was mixed with plasma (serially diluted 3X starting at 1:50) at 4 °C for 2h, then binding to HexaPro was measured by ELISA as described above.

For mAb RBD competition ELISAs to determine RBD class specificity of mAbs isolated from donors, control antibodies binding classes 1-4 (S2E12 [Class 1], P2B-2F6 [Class 2], S309 [Class 3], and CR3022 [Class 4], cloned and expressed in-house) were biotinylated using EZ-Link™ Sulfo-NHS-Biotin (Thermo Fisher Scientific) with 50-fold molar excess biotin for 48hr at 4 °C, after which free biotin was removed with 7K MWCO Zeba desalting spin columns (Thermo Fisher Scientific). Costar high binding 96-well assay plates (Corning) were coated with experimental mAbs (6 µg ml^-1^) in PBS, blocked with 2% BSA-PBS, and washed 3X with 0.1% PBS-T. RBD-SD1 was added to the plates (3 µg mL^-1^ in 50 µL PBS per well) and incubated for 1h at room temperature. The plates were washed 3X with 0.1% PBS-T and biotinylated control mAbs were added (2 x Kd in 50 uL per well). Following another wash with PBS-T, competition was determined with Streptavidin-HRP (SouthernBiotech) diluted in PBS at a ratio of 1:5000, which was detected with Ultra TMB (Millipore) as described above.

### Live-virus Neutralization assays

Live-virus neutralization assays were conducted using full-length reporter virus constructs of SARS-CoV-2 D614G (Sequence Aq. No MT020880), Delta SARS-CoV-2 B.1.617.2 (OV116969.1), Omicron BA.1 SARS-CoV-2 B.1.1.529 (EPI_ISL_6647961), Omicron BA.2.75 SARS-CoV-2 (EPI_ISL_13373170), Pangolin-CoV (EPI_ISL_410721), WIV1-CoV (KC881007.1), WIV16-CoV (KT444582.1), SARS-CoV 2003 (MK062183.1), MERS-CoV (JX869059.2), and SHC014-CoV (KC881005.1) with the nano-luciferase (nLuc) reporter gene replacing CoV ORF 7a in SARS-CoV 2003, SARS-CoV-2, WIV16-CoV, and Pangolin-CoV variants, ORF 5a in the MERS-CoV, and ORF 7 and 8 in WIV1 and SHC014-CoV, as previously published.^11–14^ For Omicron variants BA.2.12.1 SARS-CoV-2, BA.5 SARS-CoV-2, XBB1 SARS-CoV-2, XBB1.5 SARS-CoV-2,

BQ.1.1 SARS-CoV-2, and BA.2.86 SARS-CoV-2, reporter viruses were generated in a similar manner as above for SARS-CoV-2 variants, with non-synonymous amino acid mutations that differ from the ancestral Wuhan strain being incorporated into the SARS-CoV-2 spike gene based on the predominant mutations reported at the time of design for each variant.

Assays were modified from previous reports^15^ to optimize the assay system for both SARS-CoV2 and non-SARS-CoV-2 viruses. Serum dilution plates were prepared with heat-inactivated serum samples, plated at a 1:20 starting dilution, and then serially diluted 3-fold on a 96-well plate (Corning 3799) in virus growth medium (1X MEM [Gibco 11095080], 5% FBS [Hyclone SH30070.03HI] and 1% Penn-Strep [Gibco 10378016]). Monoclonal antibody dilution plates were prepared similarly to serum dilution plates, without the heat inactivation of sample, and a 1:20 starting dilution prepared from a 1 µg mL^-1^ antibody stock. Dilution plates were then transferred into the BSL3 laboratory. nLuc reporter viruses were individually diluted in the virus growth medium, added in equal volume to serum or monoclonal antibody dilution plates, and incubated for 1h at 37 °C, 5% CO_2_. The virus+serum dilutions were then transferred to duplicate columns on a 96-well black plate (Corning 3916) seeded with either Vero C1008 cells (ATCC CRL-1586) or Vero 81 (ATCC CCL-81) cells (for MERS only), and ceded at 2 x 10^4^ cells per well for a final virus dilution of 800 plaque-forming units (PFU) per well and incubated at 37 °C, 5% CO_2_. After 18-24h incubation for SARS-CoV 2003, Pangolin-CoV, WIV16-CoV, SHC014-CoV, SARS-CoV-2 D614G, Beta, Delta and 30-36h for WIV1-CoV, SARS-CoV-2 BA.1, BA.5, BA.2.12.1, BA.2.75, BA.5, XBB1, XBB1.5, and BQ1.1, the virus growth on each plate was quantified with the Promega Nano-Glo Luciferase Assay system (N1130) with a Promega GloMax Explorer (GM3500). The 50% inhibitory dilution (ID_50_) titer was defined as the serum dilution at which the observed relative light units (RLU) were reduced by 50% compared to virus+cell and virus-only control wells as determined by a Microsoft Excel macro and analyzed using GraphPad Prism 10.0.3. Additionally, all assays included monoclonal antibodies (ADG2 or S309) serving as standardized assay performance controls.

### Evaluation of mAb prophylactic efficacy in the MA10 mouse model

To test the prophylactic efficacy of SC1, SC43 and SC27, *in vivo* protection experiments were performed. For this, twelve-month-old female BALB/c mice (Envigo; 2-3 mice per group/harvest time point) were prophylactically injected 12h prior to infection with 200 µg/mouse of either mAb, isotype control mAb, or untreated. Mice were then infected intranasally with 10^3^ PFU of a mouse adapted version of SARS-CoV-2, SARS-CoV-2 MA10,^16^ or PBS. Animals were monitored daily for changes in body weight, as well as overall signs of disease. At indicated time points (2 and 4 days post-infection), mice were euthanized via isoflurane overdose, and lung tissue was harvested for viral titer analyses via plaque assay. Briefly, right caudal lung lobes were harvested into tubes containing PBS and glass beads. After homogenization, supernatant dilution series were used to infect 6-well plates containing monolayers of Vero E6 cells. Cells were overlayed with 0.8% agarose and plaques were visualized via red neutral dye 72h after infection.

### Surface plasmon resonance

Biacore CM5 sensorchips (Cytiva) were functionalized with the anti-foldon antibody MF5 using the manufacturer’s amine coupling kit. Each SARS-CoV-2 S variant (either Wuhan-Hu-1 or BA. 1) was immobilized to a CM5-MF5 sensorchip to equal RU values, washed and injected with two-fold serial dilutions of SC27 Fab. The running buffer was 1X HBS EP+ [10 mM HEPES pH 7.4, 150 mM NaCl, 3 mM EDTA, 0.005% (v/v) Surfactant P20] supplemented with 0.01% sodium azide (w/v). Response traces were double reference-subtracted and fit to a 1:1 binding model using Biacore Evaluation software.

### Biolayer interferometry ACE2 competition assay

Bio-Layer interferometry (BLI) assays were performed using an 8-channel Octet RED96e instrument (ForteBio). MF5 antibody was immobilized onto anti-human capture (AHC) biosensor SARS-CoV-2 spike was immobilized to the AHC-MF5 tips, then dipped into a solution containing 100 nM SC27 Fab until the response reached a plateau. Tips were then dipped into solutions containing 500, 250 and 125 nM ACE2. As a control, we performed the same experiment but substituted running buffer (1X HBS EP+) instead of SC27 Fab.

### Cryo-electron microscopy of SARS-CoV-2 spike (BA.1) in complex with SC27 Fab

A few minutes prior to freezing, purified SARS-CoV-2 spike (BA.1) solution was supplemented with Amphipol A8-35 a final concentration of 0.01% (w/v), then mixed with SC27 Fab at a molar ratio of 1 to 3.3 (spike trimers to Fab). The final concentration of spike in the mixture was approximately 3 mg mL^-1^. The resulting solution was incubated at room temperature for 1 min, then deposited onto glow-discharged gold grids (UltrAUfoil 1.2/1.3). Grids were plunge frozen in liquid ethane using a Mark 4 Virtobot, then loaded into a Glacios transmission electron microscope (ThermoFisher) operating at 200 kV. The microscope was equipped with a Falcon 4 detector and the pixel size was 0.94 Å. Frames were recorded in EER format. Motion correction, contrast transfer function estimation, and particle picking were performed in cryoSPARC Live, followed by 2D classification, ab initio reconstruction, and 3D refinement (heterogeneous, homogeneous, and non-uniform) in cryoSPARC v4. Masks were created in ChimeraX, then used for local refinement and 3D classification in cryoSPARC v4. Model building was performed iteratively using ISOLDE, Coot and Phenix.

### Deep mutational scanning (DMS)

Methods for the lentiviral pseudotyped DMS system were previously described by Dadonaite et al.^17^ A repo for the DMS data analysis can be found at https://github.com/dms-vep/SARS-CoV-2_XBB.1.5_RBD_DMS_SC27/. Interactive plots can be found through the documentation page at https://dms-vep.github.io/SARS-CoV-2_XBB.1.5_RBD_DMS_SC27. Complete heatmaps and line plots for Figure S4 can be found in a summary plot here: https://dms-vep.github.io/SARS-CoV-2_XBB.1.5_RBD_DMS_SC27/htmls/summary_overlaid.html.

### Statistics

GraphPad Prism version 10.0.3 (GraphPad Software Inc., La Jolla, CA, USA) was used to perform statistical analyses. Non-parametric Mann–Whitney U test and analysis of variance on ranks (Kruskal– Wallis H test) were used to determine the statistical significance of population means between two or more groups, respectively. Statistical differences in MA10 mouse modeling were tested using a one-way ANOVA with Dunnett’s multiple comparisons test, comparing every group with the mock-challenge lung titers. For plasma IgG repertoire diversity measurements, D80 is defined as the number of lineages that comprise the top 80% of the repertoire by abundance (XIC).

